# Interspecies recombination has driven the macroevolution of cassava mosaic begomoviruses

**DOI:** 10.1101/2021.04.05.438541

**Authors:** Alvin Crespo-Bellido, J. Steen Hoyer, Divya Dubey, Ronica B. Jeannot, Siobain Duffy

**Author notes:** Address correspondence to Siobain Duffy.

## Abstract

Begomoviruses (family *Geminiviridae*, genus *Begomovirus*) significantly hamper crop production and threaten food security around the world. The frequent emergence of new begomovirus genotypes is facilitated by high mutation frequencies and the propensity to recombine and reassort. Homologous recombination has been especially implicated in the emergence of novel cassava mosaic begomovirus (CMB) genotypes, which cause cassava mosaic disease (CMD). Cassava (*Manihot esculenta*) is a staple food crop throughout Africa, and an important industrial crop in Asia, two continents where production is severely constrained by CMD. The CMD species complex is comprised of 11 bipartite begomovirus species with ample distribution throughout Africa and the Indian subcontinent. While recombination is regarded as a frequent occurrence for CMBs, a revised, systematic assessment of recombination and its impact on CMB phylogeny is currently lacking. We assembled datasets of all publicly available, full-length DNA-A (n=880) and DNA-B (n=369) nucleotide sequences from the 11 recognized CMB species. Phylogenetic networks and complementary recombination detection methods revealed extensive recombination among the CMB sequences. Six out of the eleven species have descended from unique interspecies recombination events. Estimates of recombination and mutation rates revealed that all species experience mutation more frequently than recombination, but measures of population divergence indicate that recombination is largely responsible for the genetic differences between species. Our results support that recombination has significantly impacted the CMB phylogeny and is driving speciation in the CMD species complex.

**IMPORTANCE:** Cassava mosaic disease (CMD) is a significant threat to cassava production throughout Africa and Asia. CMD is caused by a complex comprised of 11 recognized virus species exhibiting accelerated rates of evolution, driven by high frequencies of mutation and genetic exchange. Here, we present a systematic analysis of the contribution of genetic exchange to cassava mosaic virus diversity. Most of these species emerged as a result of genetic exchange. This is the first study to report the significant impact of genetic exchange on speciation in a group of viruses.

## INTRODUCTION

Viruses in the *Geminiviridae* family are major constraints to agricultural crop production and pose serious threats to global food security, especially those in the genus *Begomovirus* (1). Begomoviruses are dicot-infecting, whitefly-transmitted pathogens that severely limit many economically important crops in tropical and subtropical regions around the world (2). Begomovirus genomes consist of either one (monopartite) or two (bipartite) circular single-stranded DNA (ssDNA) genetic segments, each independently encapsidated in twinned, quasi-icosohedral particles (3). There are 424 established begomovirus species in the 2019 International Committee on the Taxonomy of Viruses (ICTV) master species list, the largest number of species for any virus genus. The frequent emergence of begomovirus genotypes and persistence of begomovirus disease epidemics is facilitated by increased agricultural trade of infected plant materials, the spread of polyphagous whitefly vector biotypes (1, 4, 5), and the accelerated rate of begomovirus evolution that stems from the vast amount of genetic diversity and the consequent adaptive potential found within populations (6).

Genetic diversity is generated by a combination of mutations and genetic exchange processes (i.e., recombination and reassortment). While mutations are the fundamental source of genetic variation, genetic exchange fuels diversity by combining extant mutations from distinct genomes to produce new haplotypes. Begomoviruses have high mutation frequencies (7) and substitution rates (comparable to those of RNA viruses) (8, 9) which independently enable the efficient exploration of both sequence space and adaptive landscapes in changing environmental conditions. However, recombination has also been extensively documented among begomoviruses and is implicated in the diversification of different disease complexes affecting a variety of crops (10–13). Recombination and reassortment can introduce significant variation in a single event and profoundly impact virus evolution by preventing the accumulation of deleterious mutations (14, 15) and potentially allowing access to novel phenotypes that would be difficult to attain by mutation alone. Some phenotypic modifications associated with genetic exchange in viruses include the modulation of virulence, novel strain emergence, evasion of host immunity and antiviral resistance (16, 17).Therefore, examining patterns of viral genetic exchange is critical to understanding virus evolution and can help inform the development of control strategies.

Cassava mosaic begomoviruses (CMBs) are the causative agents of cassava mosaic disease (CMD), which frequently limits crop production in this staple food for ∼800 million people around the world (18). In 2019, Africa was the leading continent in terms of cassava yield, accounting for over 63% of the 303 million tons produced, followed by Asia with 28% (http://www.fao.org/faostat/en/#data/QC/). While the general resiliency of cassava against droughts and its tolerance of poor soil conditions has led to its widespread adoption in these regions, its susceptibility to CMD presents a major biotic constraint on production in these two continents. There are 11 identified species in the CMD species complex. Nine CMB species are found in Africa: *African cassava mosaic virus* (ACMV), *African cassava mosaic Burkina Faso virus* (ACMBFV), *Cassava mosaic Madagascar virus* (CMMGV), *South African cassava mosaic virus* (SACMV), *East African cassava mosaic virus* (EACMV), *East African cassava Cameroon virus* (EACMCV), *East African cassava Kenya virus* (EACMKV), *East African cassava Malawi virus* (EACMMV), and *East African cassava mosaic Zanzibar virus* (EACMZV). Two additional CMB species, *Indian cassava mosaic virus* (ICMV) and *Sri Lankan cassava mosaic virus* (SLCMV), have been found exclusively in Asia. The African CMB species are extensively distributed throughout sub-Saharan Africa (19) and are one of the largest threats to cassava yield, accounting for up to US$2.7 billion in annual losses (20). Although initial reports placed the Asian CMBs solely in the Indian sub-continent, SLCMV has expanded its distribution in recent years from India and Sri Lanka into Cambodia, Vietnam, Thailand, and China (21–24).

CMB genomes are bipartite, comprised of two circular segments of similar size (∼2.8 kb) which are referred to as DNA-A and DNA-B. On the virion-sense strand of the ssDNA genome, DNA-A has two partially overlapping genes that encode the coat (AV1) and pre-coat (AV2) proteins. The complementary strand of DNA-A encodes the replication-associated protein (AC1), the transcriptional activator protein (AC2), a replication enhancer (AC3) and an RNA-silencing suppressor (AC4). The DNA-B segment encodes for two proteins - a nuclear shuttle protein in the virion sense (BV1) and a movement protein in the complementary sense (BC1) (25, 26). Although genetically distinct, both segments share a common region (CR) of ∼200 nucleotides that includes a stem loop structure with the conserved nonanucleotide TAATATTAC where rolling-circle replication is initiated. Additionally, the CR contains several regulatory elements including multiple copies of cis-elements known as iterons which are binding sites for the replication-associated protein (27).

Analyses from field samples have revealed that both CMB segments are frequently evolving through homologous recombination (and “recombination” is presumed to be homologous recombination in this manuscript) (28–34). Most notably, recombination contributed to the emergence of a highly virulent hybrid of ACMV and EACMV isolates known as EACMV-Uganda (EACMV-UG) that caused severe disease outbreaks in East and Central Africa in the 1990s (35, 36). Due to the frequent characterization of emergent recombinants and the fact that distinct CMBs are commonly found infecting the same plant (37–39), recombination is regarded as a widespread phenomenon that significantly impacts CMB biodiversity and evolution.

Here we present a systematic analysis of recombination and its influence on the evolution of the CMD species complex. By applying several recombination analysis tools to datasets of publicly available CMB sequences, we mapped a complex recombination history where inter-species recombination events correlated with the emergence of most (6/11) CMB species. While mutation was estimated to occur more often than recombination in all our datasets, our findings support interspecies recombination as the main driver of diversity at a macroevolutionary scale.

## RESULTS

A total of 880 full-length DNA-A sequences and 369 DNA-B sequences from the eleven established CMB species were downloaded from NCBI GenBank (Table 1). The DNA-A isolates were classified based on the begomovirus 91% nucleotide identity species demarcation threshold. Pairwise nucleotide identity comparisons (Supplemental file 1) resulted in the reassignment of four isolates previously identified as EACMV sequences to EACMCV (accessions: AY211887, AY795983, JX473582, MG250164).

**Table 1.**
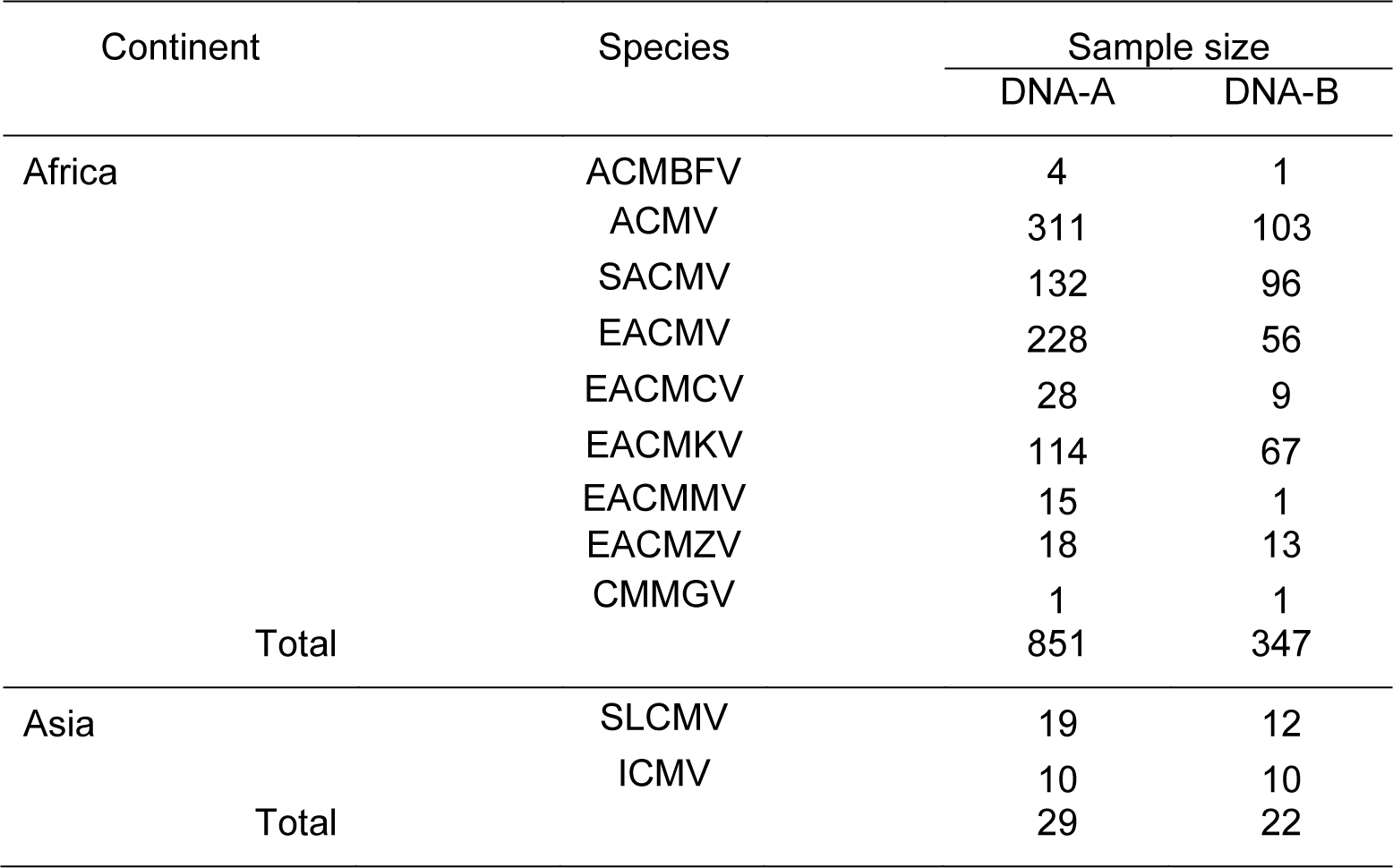
Sample sizes by designated species for DNA-A and DNA-B sequences used in this study

Because the species definition does not extend to the DNA-B segment, DNA-B sequences were identified according to their species designation in GenBank (DNA-B segments are typically classified based on DNA-A sequences isolated from the same host sample or by highest nucleotide identity to an extant DNA-B sequence when no corresponding DNA-A sequence is available). Our datasets are imbalanced with respect to genomic segment (DNA-B is less frequently sampled than DNA-A) and geography (the sample size was larger for African CMBs relative to Asian CMBs). We present results for DNA-A followed by DNA-B.

### Likely recombinant origin for 6 of 11 CMB species

Since recombination is a major contributor to begomovirus evolution, standard phylogenetic approaches cannot fully recapitulate the evolutionary history of CMBs. Therefore, we used a split-network analysis to examine evolutionary relationships within the CMB phylogeny. The network (Fig.1A) showed most sequences in tight clusters based on the 11 species. Some divergent isolates were found near the main clusters in the SACMV, EACMKV and EACMV clades, suggestive of phylogenetic conflict and, potentially, recombination causing the divergence in those sequences. Multiple edges connecting branches of SLCMV and ICMV isolates indicate complicated patterns of recombination among the Asian CMBs, consistent with previous reports (40). The highly reticulate structure of the network implies an extensive history of recombination, both within and between species.

**Figure 1.**
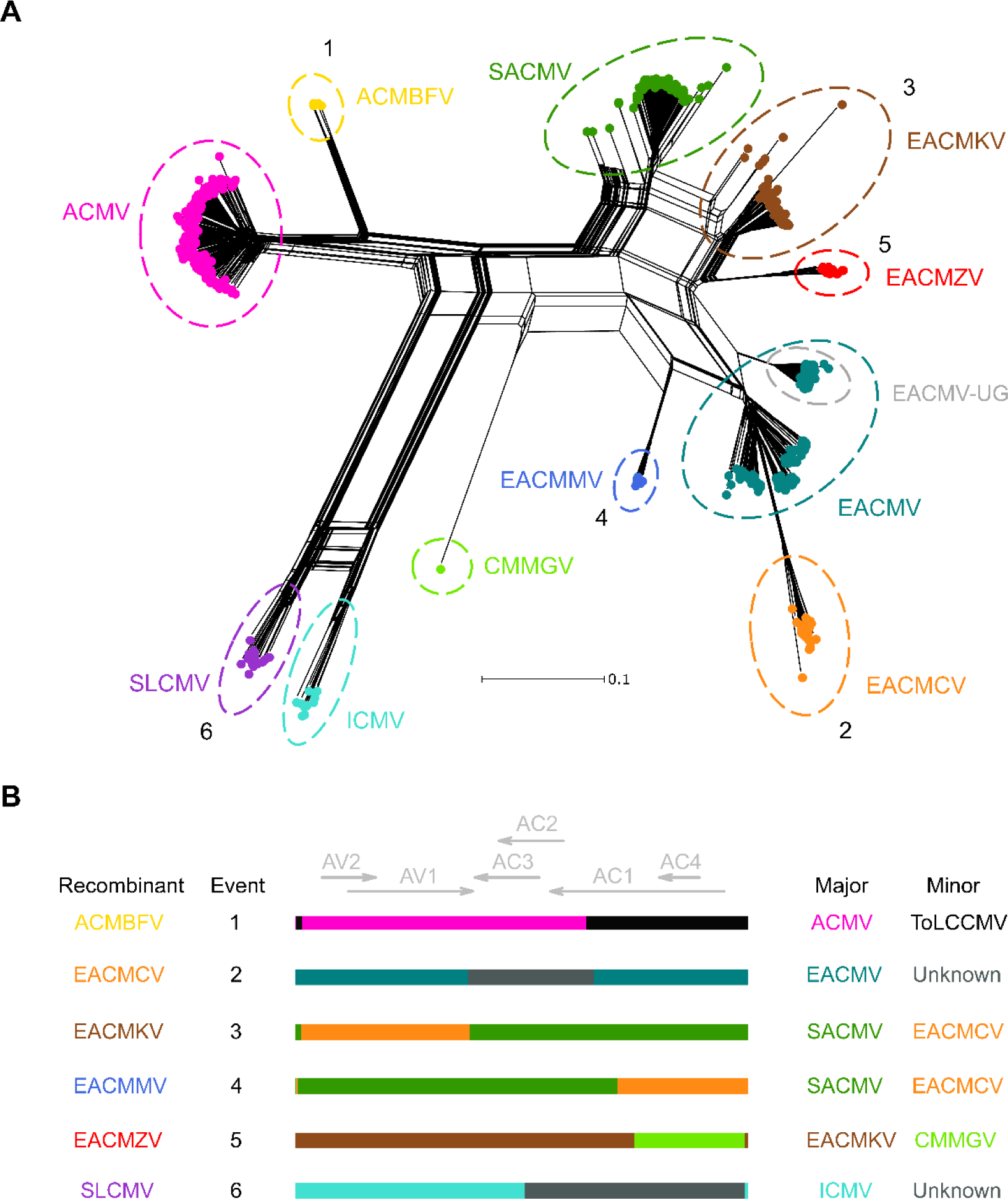
Phylogenetic network analysis for all CMB species DNA-A segments and schematic of identified macroevolutionary recombination events. (A) Neighbor-net network analysis for all CMB species DNA-A segments. Distances were transformed using a GTR + G model. The numbers correspond to the clade-wide events shared by all members of a species reported in Table 2 and depicted in 1B. Colors representing each species are: ACMV: pink, ACMBFV: yellow, SACMV: green, EACMKV: brown, EACMZV: red, EACMV: teal, EACMCV: orange, EACMMV: blue, CMMGV: spring green, ICMV: turquoise, SLCMV: purple. The descendants of a well-studied recombination event between ACMV and EACMV (EACMV-UG) are circled in gray. (B) Linearized DNA-A schematic representations of putative ancestral recombination events based on breakpoint and parental species predicted by RDP4.

**Table 2.**
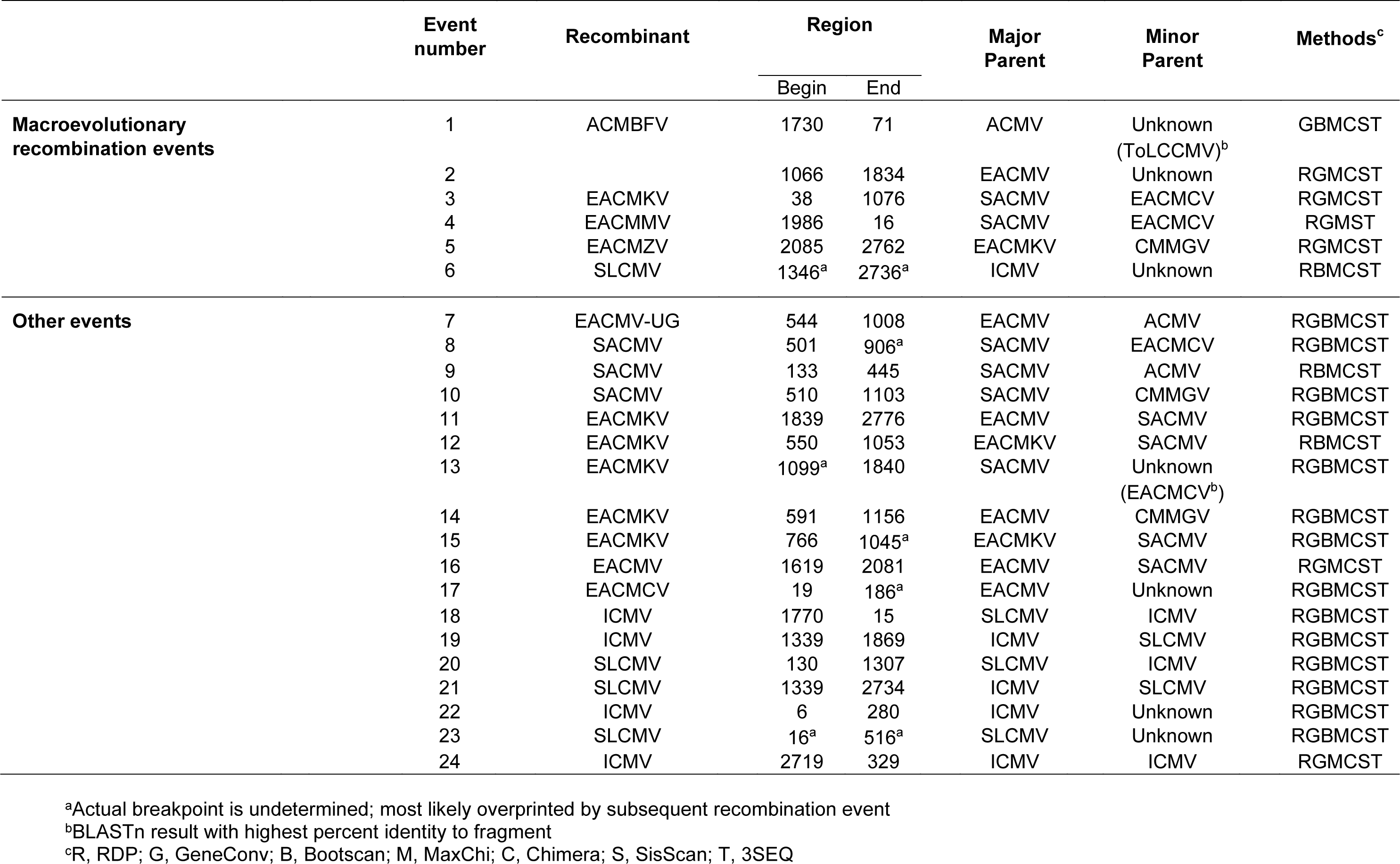
List of recombination events detected in CMB DNA-A sequences

To further explore and characterize recombination among the CMB DNA-A sequences, the all-species alignment (n=880) was analyzed using RDP4. An initial scan did not detect recombination between the Asian and African sequences. We split these sequences into two data sets (African n=851; Asian n=29) with the rationale that reducing the number of gaps in the alignments would improve accuracy of recombination detection. We performed RDP4 analysis on the two multiple alignments separately (with stringent settings, described in the Methods section) and identified a total of 24 high-confidence recombination events (Table 1): 16 for the African CMB data set and 8 for the Asian data set. Six unique events were supported in all representatives of individual species (depicted in Fig. 1B; similarity plots in Fig. A1 and A2), suggesting a recombinant origin for six out of the eleven species: ACMBFV, EACMCV, EACMKV, EACMMV, EACMZV and SLCMV. We refer to these events as ‘macroevolutionary,’ based on the hypothesis, discussed below, that the recombination events led to the original splitting of each relevant species cluster from “parent” species clusters. Most of these events have been reported previously, except for that associated with SLCMV. Similarity plots for all 24 high-confidence events using the best candidate parental sequences identified by RDP4 are presented in the appendix.

As in Tiendrébéogo et al. (32), ACMBFV was identified as a recombinant of ACMV with a recombinant fragment spanning most of the AC1 ORF, the entire AC4 ORF and a portion of the CR. Despite RDP4 choosing CMMGV as the minor parent for this event in our analysis, low nucleotide identity (<80%) within the recombinant region makes it an unlikely parental sequence (Fig. A1). BLAST analysis of the recombinant portion identified a tomato leaf curl Cameroon virus (ToLCCMV) sequence as the closest ‘relative’ currently in GenBank, which is consistent with the previous report. The EACMCV and EACMZV macroevolutionary recombination events (events 2 and 5; Fig. A1) corroborate results from previous recombination analyses where they were characterized as recombinants (29, 30). No significant virus donor was identified for the EACMCV recombinant fragment, but its major parent was likely EACMV. EACMMV has been described as an EACMV-like recombinant (28, 41), yet RDP4 suggested SACMV as the most likely major parent in our analysis. The conflict between these results is most likely due to the very high degree of similarity between the regions covering AC3, AC2 and the 3’ end of AC1 in SACMV and EACMV (Fig. 2A), which suggests a shared evolutionary history for that region among the two species. Since high similarity can confound recombination detection, it becomes hard to unambiguously detect correct breakpoints and potential parental sequences. However, analysis with similarity plots showing a drop-off in similarity at one of the boundaries of this region between EACMV and EACMMV, points to SACMV being the more likely major parental species (scenario 1 illustrated in Fig. 2B). The high-sequence-similarity region also affects candidate parent sequence identification for EACMKV (Fig. 2C, discussed below).

**Figure 2.**
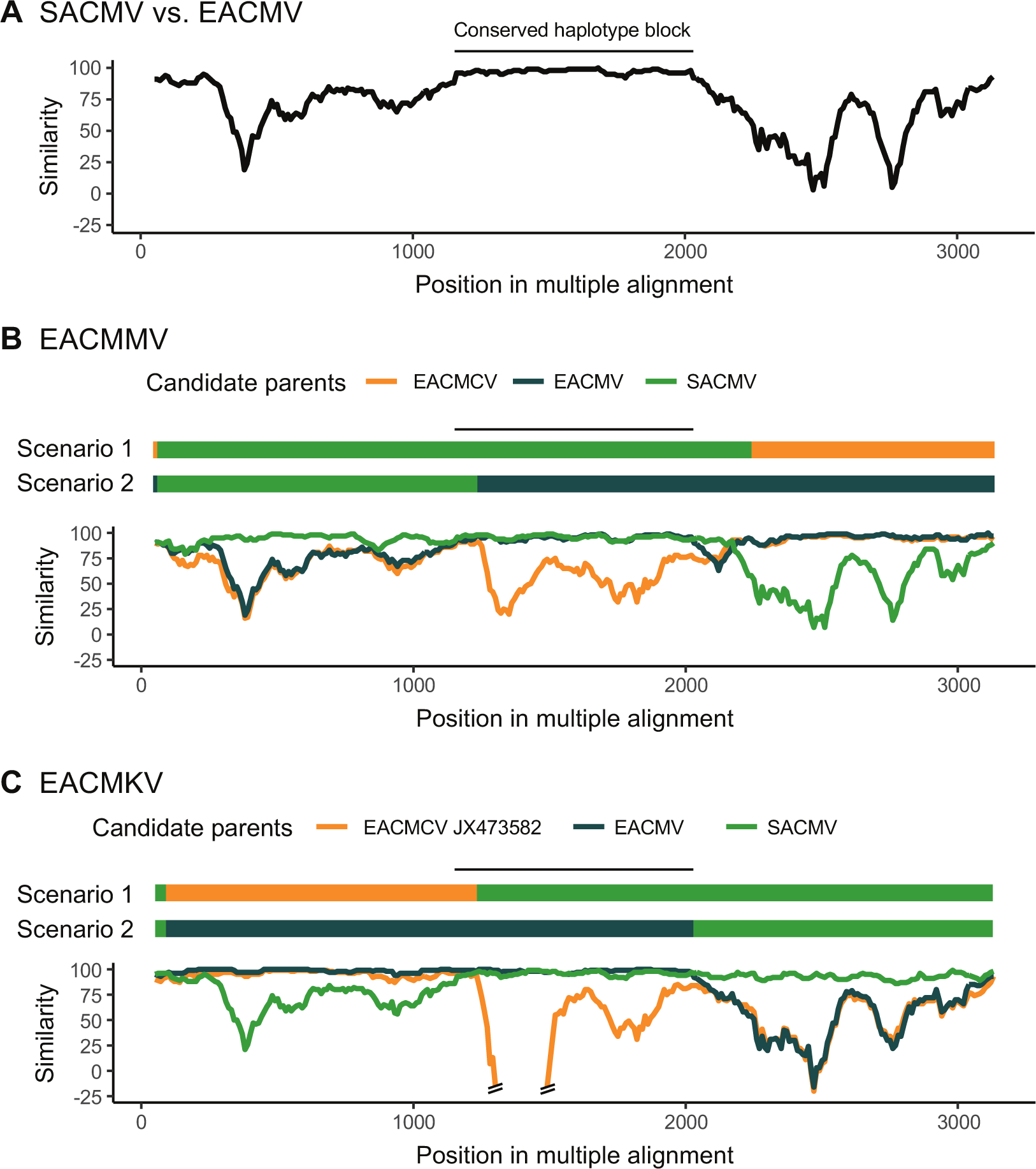
Similarity plots showing similarity between EACMV and SACMV (A) and alternative recombinant origins for the EACMMV (B) and EACMKV (C) DNA-A segments. Plots are comparing the similarity of 50% consensus sequences of the plotted species (except for 2C where best candidate EACMCV_JX473582 was used) against a 50% consensus sequence of the query species. The Kimura-2-Parameter model of sequence evolution was used to correct distances and a sliding window size of 100nt with a step size of 10nt was used. Above each plot, Scenario 1 represents the macroevolutionary events associated with the queried species based on RDP4 results while Scenario 2 represents the plausible alternative recombinant origin. The black line above the schematics depicts the conserved region between EACMV and SACMV sequences, derived from panel 2A.

Curiously, the single available CMMGV sequence, which has been previously characterized as recombinant (33), did not display any putative recombinant regions within its genome. It was reported that CMMGV had minor fragments donated by both SACMV and EACMZV-like sequences. However, a close examination using the distance plot and phylogenetic tree construction tools in RDP4 revealed that only one SACMV sequence (the minor parent identified by Harimalala et al., and the first SACMV isolate ever fully sequenced (42)); accession number: AF155806) had high similarity in the AV1 recombinant region with CMMGV (event 10; Fig. A4), whereas all other SACMV genomes did not. As a result, it seems more plausible that CMMGV acted as donor to that single SACMV isolate. In the case of the second minor fragment, our RDP4 analysis suggested that CMMGV was the donor virus and EACMZV the recipient (event 5; Fig. A1), contrary to what was argued previously (33). At the moment, we cannot distinguish the direction in which the fragment was donated so there is no definitive evidence as to whether CMMGV is a recombinant species.

The results showed frequent recombination between ICMV and SLCMV, which made it difficult to resolve the recombination profiles within the Asian CMB data set.

This issue is suggested by the statistically undetermined breakpoints in the SLCMV species-wide event involving ICMV and an unknown minor parent (event 6; Fig. A6), which points to likely overprinting by subsequent recombination events. Only 16 out of 19 SLCMV sequences were predicted to be descendants of this event. However, the three remaining SLCMV isolates (accessions: AJ314737, KP455484, and AJ890226) showed evidence of a similar event between ICMV and an SLCMV-like isolate with almost the same breakpoints (event 21; Fig. A7). Altogether, we interpret these results as evidence for a recombinant origin for SLCMV.

### Other high confidence DNA-A events confirm previously described recombinants

In addition to the six macroevolutionary events, 18 other events were detected in the DNA-A datasets (Table 2). Among these events, the most well-represented event was that of the famous EACMV-UG recombinant (event 7; Fig. A3), found in 97 of the 228 EACMV sequences. Of the 10 other non-macroevolutionary events in the African data set, most were associated with either EACMKV or SACMV as the recombinants (5 and 3 events, respectively). Recombinants with evidence of events 8, 9, 12, 13, 14, 15 and 17 (Fig. A3-A5) were collected in one of the most comprehensive CMB sampling studies to date, which took place in Madagascar (34). The EACMKV isolate in event 13 (accession: KJ888083) presented an interesting case as it was classified as EACMKV by having 91.02% nucleotide identity to only one other EACMKV isolate (accession: KJ888079; Supplemental file 1), suggesting the event caused just enough divergence to where the sequence narrowly satisfies the criterion to be classified as EACMKV. A BLAST analysis revealed an EACMCV isolate from Madagascar (accession: KJ888077) as a highly similar recombinant donor, which was not identified by RDP4 as a parent despite being present in the dataset.

Recombination event 11 could conflict with the recombinant origin for all other EACMKVs, as all EACMKV sequences match both this and the profile suggested in event 3 (Fig. 2B). We maintain that event 3 is the more likely origin of the EACMKV species on the basis that it was detected in all EACMKV sequences in our analysis. However, the alternative recombinant origin where an EACMV sequence acts as the major parent is consistent with the first characterization of an EACMKV isolate (31) and remains a possibility. This highlights once more the challenge in characterizing events involving SACMV and EACMV-like sequences due to their region of high similarity (Fig. 2A). No evidence of recombination was found among EACMZV and EACMMV isolates.

Despite having a smaller pool of sequences, 4 genetic exchanges between SLCMV and ICMV were identified in the Asian data set. Three of those events (6, 19 and 21; Fig. A6-A7) had breakpoints in the region of overlap between AC2 and AC3, suggesting a potential hotspot of recombination between these species.

### Mutation occurs more frequently than recombination within the DNA-A segment of all CMB species

We estimated nucleotide diversity (π) within all species (except for CMMGV and ACMBFV, which each had fewer than 5 sequences, Table 1) as a measure of standing genetic diversity. Nucleotide diversity for all species was within the same order of magnitude and ranged from 0.012 (for EACMMV) to 0.074 (for ICMV, Table 3). No associations were observed between diversity and sample size.

**Table 3.**
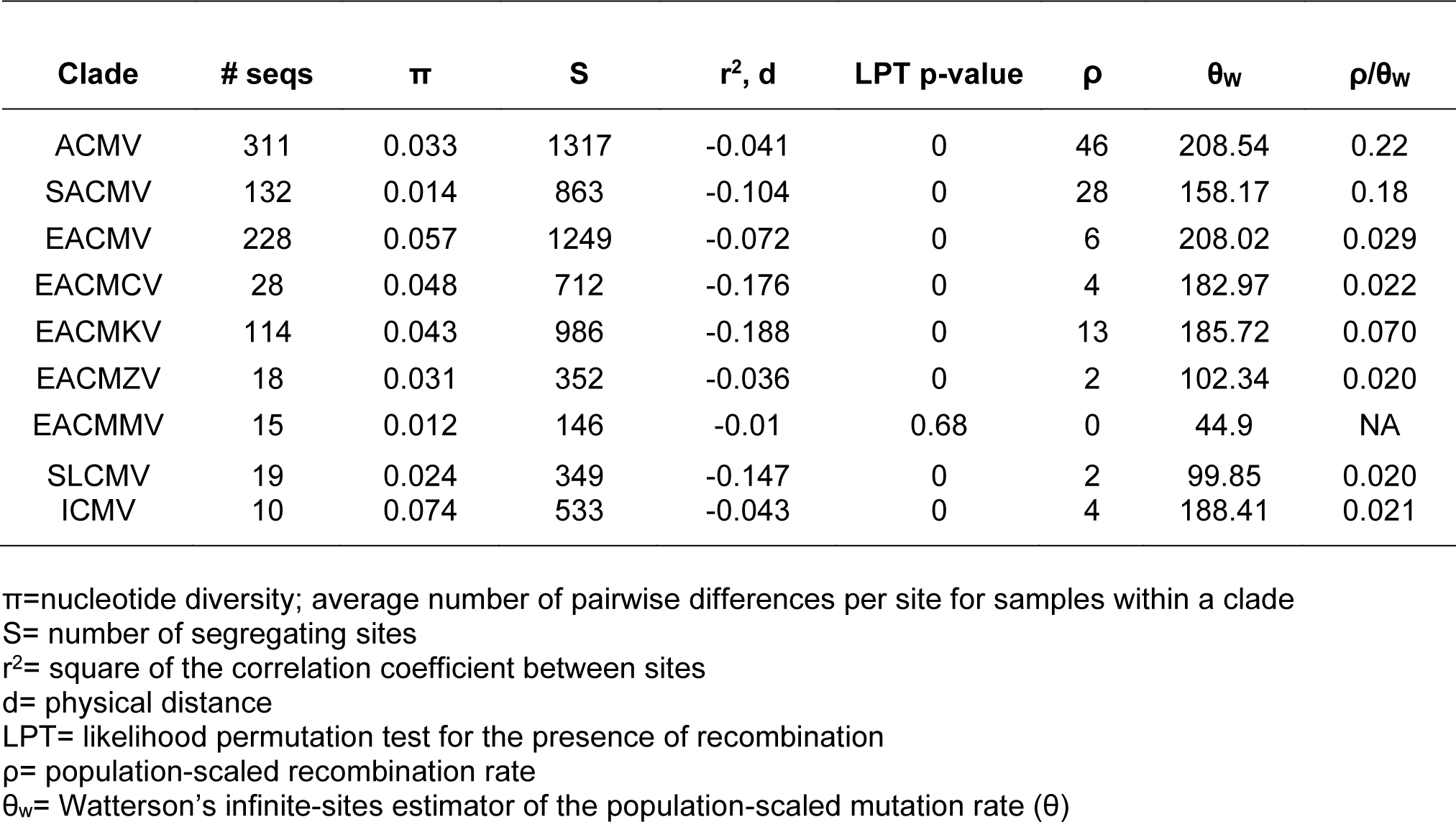
Descriptive statistics of CMB DNA-A species’ diversity and the contributions of mutation and recombination to that diversity

Additionally, we estimated per-generation, population-scaled rates of recombination (ρ) and mutation (θ) to assess the frequency of recombination within each species relative to mutation (ρ/θ). We further tested for the presence of recombination by calculating correlations between estimates of linkage disequilibrium (r^2^) and physical distance (d) and used a likelihood permutation test (LPT) of recombination (Table 3) with LDhat (43).

The correlation between r^2^ and d was negative across all datasets, consistent with the expectation of linkage disequilibrium decay as distance is increased in the presence of recombination. The LPT indicated recombination in all species except EACMMV, which was consistent with the ρ=0 estimate for that species. Across all populations, mutation was the dominant evolutionary mechanism in terms of frequency when compared to recombination, as displayed by <1 values of the ρ/θ ratio (typically < 0.03). Interestingly, the highest ρ/θ value was observed for ACMV (0.22), which was involved in three interspecies recombination events detected thus far (events 1, 7 and 9), but none within the species. SACMV and EACMKV were the other two clades with higher contributions of recombination, which were also the two most featured species in our RDP4 results for the African CMB sequence alignment.

### Sequence divergence between DNA-A recombinant species and their hypothesized major parents suggests interspecies recombination as the major contributor to phylogenetic divergence

The average number of pairwise nucleotide differences per site within species (π) and between recombinant and predicted major parental species (D_XY_) were estimated in sliding windows to assess the effect of the macroevolutionary recombination events on phylogenetic divergence (Fig. 3). In every comparison, there was a pronounced increase over the genome-wide average of D_XY_ in regions associated with macroevolutionary recombination events. This suggests appreciably different evolutionary histories in those regions compared to the rest of the genome, which supports species-wide recombination events as drivers of greater divergence than mutational and other minor recombination events. A noticeable peak in D_XY_ within the CR and 5’ end of AV2 in the EACMCV-EACMV comparison was observed (Fig. 3). This region was detectably recombinant in one EACMCV sequence (event 17, accession: KJ888049). Close examination of the alignment in this region suggested that 23 of the remaining 27 EACMCV sequences may have an undetected recombination event in this region, but likely with different breakpoints from event 17.

**Figure 3.**
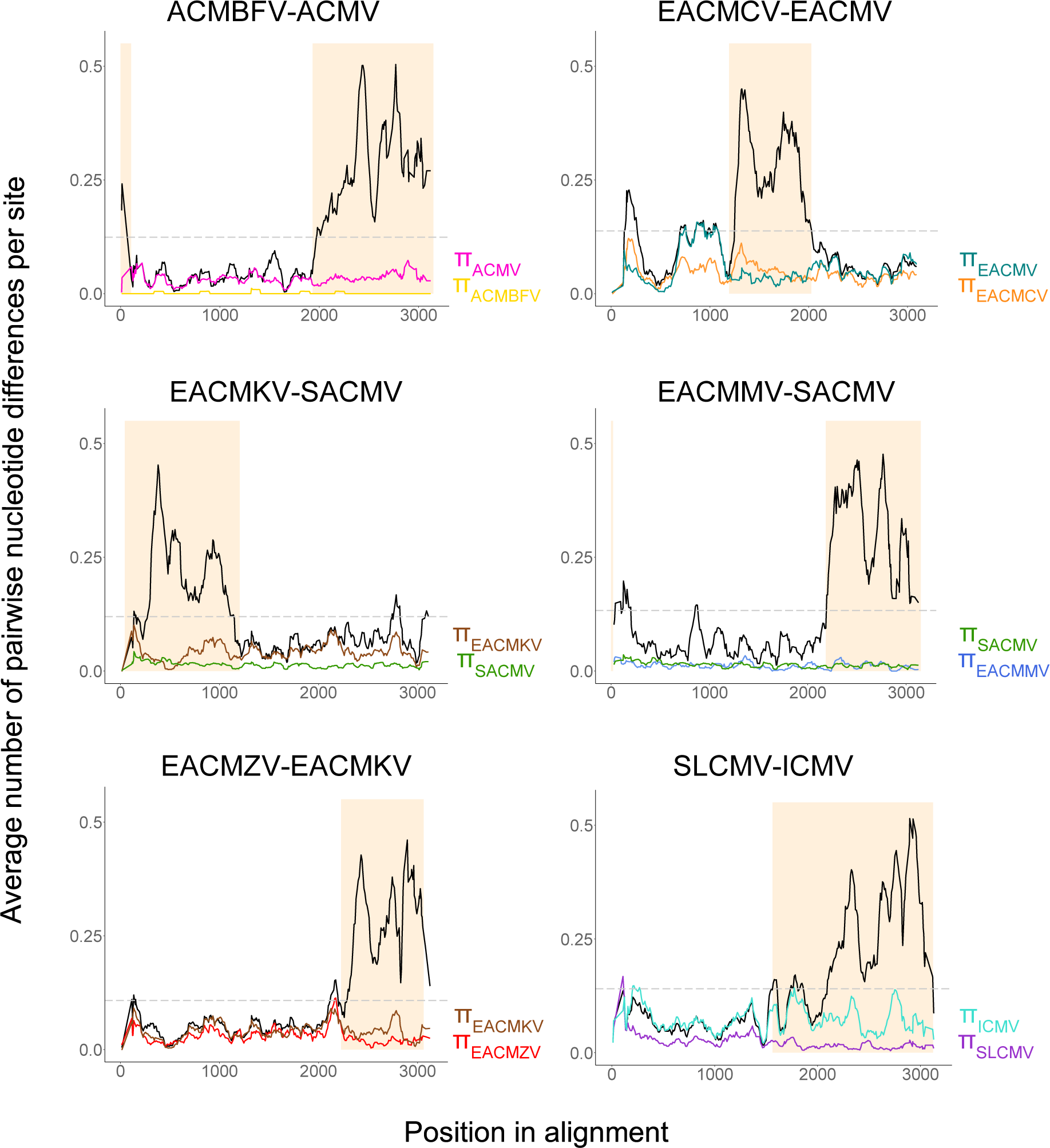
Sliding window plots of nucleotide diversity (π) and population divergence (DXY) for CMB DNA-A recombinant-major parental clade combinations. DXY between species is plotted in black and the grey dashed line represents the DXY average between the compared datasets. The shaded area depicts the predicted recombinant fragment based on RDP4 analyses.

All samples with evidence of the undetected event and event 17 were sampled in West Africa, Comoros or Madagascar while the three EACMCV isolates without a recombination event in that part of the genome were sampled in East Africa. This supports the hypothesis that EACMCV originated in East Africa and acquired a second recombinant fragment in the West African isolates (41), and it is possible that the West African genotype has now been introduced to the Comoros and Madagascar. The uncharacterized event and event 17 clearly have contributed to the divergence within EACMCV (as evidenced in a spike in EACMCV nucleotide diversity; Fig. 3) and between EACMV and EACMCV. Similarly, a downstream increase in D_XY_ and π for EACMV within the AV1 3’ end was observed, corresponding to the region of the EACMV-Ug recombination event (event 7). While these are examples of how small recombination events have contributed to the phylogenetic divergence between species, our results show that the larger, ancestral inter-species recombination events are the driving force behind evolutionary divergence at the CMB species level.

### Fewer high-confidence DNA-B recombination events

Due to the high levels of divergence between DNA-B isolates, an all-“species” DNA-B alignment was difficult to construct. Therefore, we split the DNA-B sequences into three broad groups: EACMV-like (EACMV, SACMV, EACMKV, EACMMV, EACMZV, EACMCV) + CMMGV (n=243), ICMV-SLCMV (n=22) and ACMV-ACMBFV (n=104), and conducted phylogenetic network (Fig. 4A) and RDP4 analyses (Table 4) on each group separately. For the EACMV-like group, we observe a network with sporadic reticulations indicating some recombination. The DNA-B sequences from most species do not form monophyletic clades, with isolates from EACMV, EACMKV and SACMV spread out around the network. Isolates from EACMCV, which have been reported as clearly distinct from the rest of the EACMV-like DNA-B segments (44), are separated from the center of the network by long branches, indicating large genetic distances between them and the rest of the EACMV-like DNA-B segments. Similarly, a long branch separates CMMGV from all other clusters.

**Figure 4.**
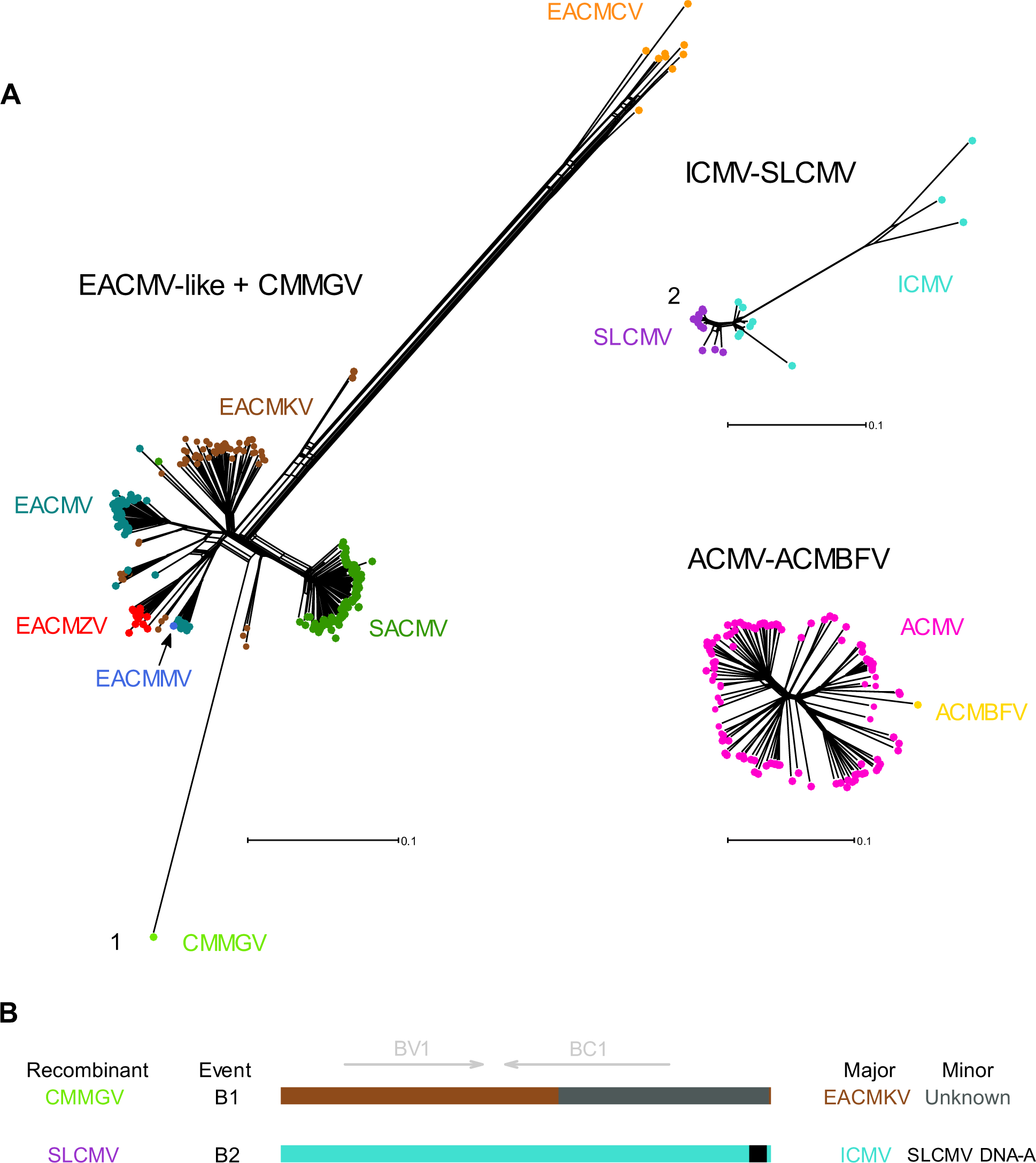
Phylogenetic network analysis for all CMB species DNA-B segments and schematic of identified clade-forming recombination events. (A) Neighbor-net network analysis for groups of CMB DNA-B segments. Distances were transformed using a GTR + G model. The numbers correspond to the events reported in Table 2 and depicted in 4B. (B) Linearized DNA-B schematic representations of putative species-spanning recombination events based on breakpoint and parental species predicted by RDP4.

**Table 4.**
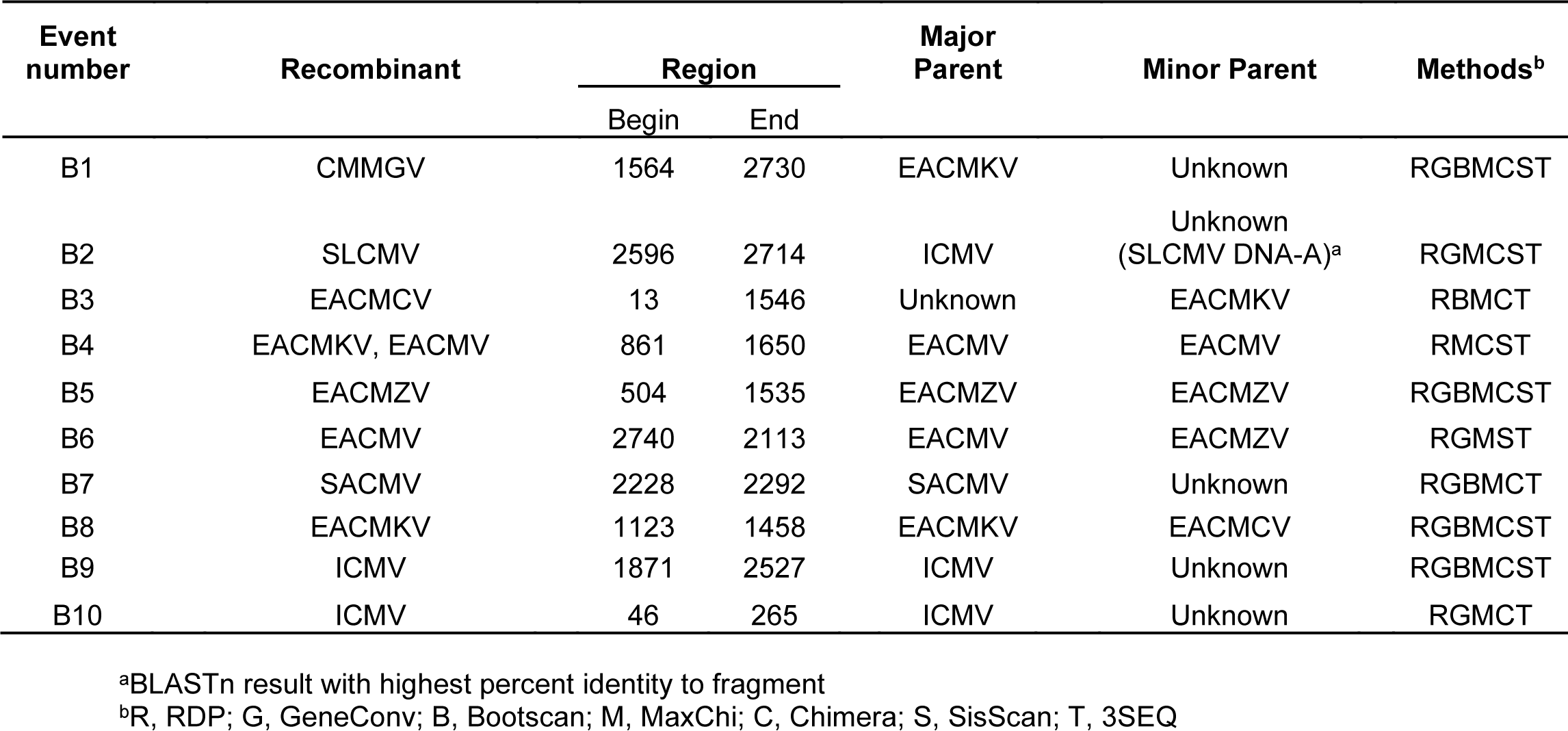
List of recombination events detected in CMB DNA-B sequences

The ICMV-SLCMV network is more compact than the EACMV-like group, signifying a higher degree of genetic similarity between all isolates. All the SLCMVs are closely related to one another, and the branches for both SLCMV and ICMV isolates show some reticulation. The ACMV-ACMBFV sequences are also genetically very similar and have the least reticulation of the three networks (Fig. 4A).

A total of 10 recombination events were identified in the DNA-B data sets: 7 events in the EACMV-like group and 3 events in the ICMV-SLCMV group (designated B1-B10, Table 4). No events were detected in the ACMV-ACMBFV sequences. Of the 10 events, 2 could be considered as ancestral clade-founding events (Fig. 4B). We refer to these events as ‘clade-founding’ rather than ‘macroevolutionary’ to emphasize that this classification is distinct from the DNA-A-based species definition. Event B1 is associated with CMMGV (Fig. A9), which had an EACMKV isolate as a closely related major parent and a recombinant fragment from an unknown virus that spanned most of BC1 and the 5’ portion of the CR. This event was previously reported (33). Event B2 was associated with SLCMV (Fig. A9), where all 12 sequences had evidence of the event. In this event, ICMV was observed as major parent with a fragment in the 5’-CR from an unknown parent. From a BLAST analysis, we identified that the fragment most likely originated from an SLCMV DNA-A sequence. This event has been described before and is believed to explain the evolution of SLCMV from a putative monopartite begomovirus, where an SLCMV-like sequence “captured” an ICMV DNA-B segment by donating the Rep-binding iteron sequences necessary for replication (45).

In addition to these two well-supported clade-forming events, there are several other recombination events that may have had a similar impact. Seven out of the 9 EACMCV DNA-B sequences show evidence of event B3 (Fig. A9), and it is plausible that this recombination event defined the common ancestor of all 9 EACMCV DNA-B sequences. However, the lack of statistical support in the other two sequences prevents us from calling it an ancestral recombination event for all EACMCV DNA-B isolates.

Similarly, event B4 (Fig. A10) is observed in 8 different sequences classified as either EACMKV or EACMV, which suggests recombination was the mechanism of emergence for this small circulating clade. Event B8 (Fig. A9) is another example of an event that possibly led to the emergence of a small clade, identified in two EACMKV isolates (accessions: JF909228 and JF909227) collected in the Seychelles archipelago (46).

No recombination meeting our 5-out-of-7-methods RDP4 threshold was found in the ACMV-ACMBFV DNA-B group. However, one event in the single ACMBFV DNA-B sequence available was detected by 4 methods. This event was not detected in the original report of ACMBFV (32) but similarity plots (Fig. A8) provide additional evidence of recombination. It seems likely that ACMBFV DNA-A “captured” an ACMV DNA-B segment via recombination, creating an ACMBFV DNA-B segment with a compatible replication-associated protein binding site (Fig. A8), similar to the scenario proposed for SLCMV DNA-B (45). An alternative possibility is that an ACMV DNA-B molecule recombined directly with a different virus segment (based on best BLAST hit, potentially a relative of tomato leaf curl Nigeria virus – accession: FJ685621).

### Mutation occurs more frequently than recombination within CMB DNA-B groups

Nucleotide diversity and rates of mutation and recombination were estimated for ACMV-ACMBFV, EACMCV and ICMV-SLCMV groupings (Table 5). The high degree of similarity within these groups justifies them being defined as individual “populations” for these analyses. The other species were not included in this analysis because of our inability to define meaningful populations. Nucleotide diversity estimates for the ACMV-ACMBFV DNA-B cluster were higher (0.067) than for ACMV DNA-A (Table 3, 0.033).

**Table 5.** Descriptive statistics of CMB DNA-B groups’ genetic diversity and the contributions of mutation and recombination to that diversity.

| Clade | # seqs | $\pi$ | S | $r^2$, d | LPT p-value | $\rho$ | $\theta_w$ | $\rho/\theta_w$ |
| --- | --- | --- | --- | --- | --- | --- | --- | --- |
| ACMV-ACMBFV | 104 | 0.067 | 1335 | -0.009 | 0.003 | 28 | 193.31 | 0.145 |
| EACMCV | 9 | 0.088 | 711 | -0.005 | 0 | 3 | 261.60 | 0.011 |
| ICMV-SLCMV | 22 | 0.062 | 881 | -0.036 | 0 | 2 | 241.68 | 0.008 |

$\pi$ = nucleotide diversity; average number of pairwise differences per site for samples within a clade
S = number of segregating sites
$r^2$ = square of the correlation coefficient between sites
d = physical distance
LPT = likelihood permutation test for the presence of recombination
$\rho$ = population-scaled recombination rate
$\theta_w$ = Watterson's infinite-sites estimator of the population-scaled mutation rate ( $\theta$ )

The same was observed for the EACMCV DNA-A and DNA-B segments (0.048 and 0.088, respectively). However, we estimate a slightly higher standing genetic variation in the ICMV DNA-A sequences (0.074) than in the ICMV-SLCMV DNA-B group (0.062), indicating comparable levels of variability among the examined sequences.

We found evidence of linkage disequilibrium decay in all three datasets using the r^2^ measure, and the LPT indicated the presence of recombination in all groups. The ρ/θ ratio ranged from 0.008-0.145, showing that mutation is much more frequent than recombination for DNA-B. Overall these within-group results for DNA-B (Table 5) were very similar to those for DNA-A (Table 3).

## DISCUSSION

Recombination is an important and pervasive mechanism that contributes significantly to plant virus evolution (47) and is broadly documented among begomovirus species (10–13, 16). Our updated recombination profile of all sampled CMB full genomic segments to date reveals widespread intra- and inter-species recombination. A variety of complementary recombination analyses indicate that the majority of CMB species (6/11) have a recombinant origin. For the first time, we show a recombinant origin of SLCMV DNA-A, which likely descended from genetic exchange between an ICMV-like isolate and an unidentified begomovirus DNA-A segment.

Surprisingly our analysis did not support a recombinant origin of the single isolate of CMMGV, though it had been considered previously to be the product of genetic exchange between major parent distantly related to CMBs and minor parents SACMV and EACMZV (33). Instead, our analyses consider CMMGV to be a parental virus, contributing to the creation of EACMZV and an SACMV recombinant (42). Although no macroevolutionary signals of recombination were detected in SACMV, ACMV, CMMGV, EACMV and ICMV, it is possible that events associated with their emergence occurred so long ago that the distinguishing patterns of polymorphism created by recombination have been erased by subsequent mutations and cannot be detected. In the case of SACMV, an argument has been made for it having a recombinant origin based on molecular analyses and phylogenetic incongruencies observed in different parts of the genome where the AV1 ORF and CR resemble tomato yellow leaf curl virus isolates, the AC2, AC3 and AC1 3’ end are closely related to EACMV, and the 5’ end of AC1/AC4 ORF portion seems to have a distinct evolutionary history (41, 42, 48, 49). Regardless of the undetectable contribution of recombination to all CMB DNA-As, we have strong evidence for recombination leading to speciation in the majority of currently defined CMB species.

Although parentage cannot be definitively established in some cases, fragments derived from EACMKV and SACMV lineages seem to have a high propensity for inter-species recombination (34). We also observe frequent recombination between both Asian CMB species, which supports past reporting of ICMV and SLCMV as a recombinogenic pair (50). Unsurprisingly, no recombination was detected between isolates originating in Asia and those from Africa. At this moment, there are no reports of Asian CMBs infecting cassava crops in Africa and there is only one study where an African CMB (i.e., EACMZV) has been sampled in cassava crops in Asia, specifically in the West Asian country of Oman (51), where ICMV and SLCMV have never been identified. While we lack experimental or field evidence that Asian and African CMB species have the capability to recombine and produce viable viral progeny, it is probable that these viruses have had limited opportunities to coinfect the same host plant.

However, ACMV isolates have been recovered in cotton crops in the South Asian country of Pakistan and ACMV recombinant fragments have been detected in cotton-infecting begomoviruses even though cassava is not cultivated there (52). This suggests an exchange of CMB species between Africa and Southern Asia and hints at the role of alternative plant hosts on the emergence of inter-species recombinant begomoviruses (53). Continuous CMB surveillance efforts are needed to ensure endemic viral species are not spread across continents via international trade and are not given the opportunity to spread and potentially recombine with native begomoviruses. Countries heavily involved in agricultural trade such as Oman should receive special attention as they can become a sink for divergent begomovirus species and potentially a hub for the emergence of novel recombinant begomoviruses (54).

### While mutation is more frequent, retained recombination events are more significant

Estimates of the DNA-A relative rates of recombination and mutation (ρ/θ) show that mutations occur more often than recombination within all the analyzed clades, which is consistent with previous analysis based on the Rep and CP genes of other begomovirus species (55). Notably, while no recombination was readily detected in any of the ACMV sequences with RDP4, the LPT detected a signal of recombination, and our ACMV DNA-A dataset had the highest frequency of genetic exchange relative to mutation. Since recombination signals were detected within the ACMV sequences, we interpret these results collectively to mean most ACMV recombination is intra-specific (illustrated in Fig. 1), which is difficult to detect with the methods used by RDP4. The lack of ACMV recombinants involving other CMB species, which has been mentioned in the literature (20, 41), might be explained by potential genome incompatibility and selection against mosaic sequences where donor fragments from divergent CMBs could disrupt intra-genomic interactions and gene coadaptation (56, 57).

Mutation is clearly the more frequent process when compared to recombination within all CMB species, confirming the conclusions of previous studies which show that the genetic diversity in begomovirus populations is predominantly shaped by mutation (10, 55). However, the relative contributions of these processes to the evolution of CMBs is not necessarily a function of their frequency. The D_XY_ sliding-window plots reveal that single recombination events are correlated with most of the divergence between putative parental species and their recombinant progeny species (Fig. 3). We conclude that the relatively higher rates of mutation relative to recombination on a microevolutionary scale are not reflective of the influence of recombination at the macroevolutionary scale, where inter-species recombination is the driving force behind the emergence of the majority of CMB species. While reports of begomovirus species that have emerged through recombination are common (58–62), this is the first time a systematic analysis of recombination and its contribution to species diversity is performed within all known species of a begomovirus disease complex.

Although “speciation” does not directly apply to DNA-B sequences it is clear that intersegment recombination has played a significant role in the evolution of viruses such as SLCMV and, potentially, ACMBFV. Indeed, the trans-replicational capture of divergent DNA-B segments/satellite molecules by DNA-A sequences via recombination involving Rep-binding sites is a documented mechanism that has led to new associations resulting in different disease phenotypes (45, 63, 64). Additionally, it represents a plausible explanation for the potential evolutionary transition to bipartite begomoviruses from monopartite ancestors (44). Ultimately, this phenomenon has and continues to contribute to the genomic modularity of begomoviruses, which in turn can influence their evolvability.

The phylogenetic networks among the DNA-B groups (Fig. 4) were comparatively less reticulated than the DNA-A network (Fig. 1), and fewer high-confidence recombination events and recombinants were detected. Despite these comparisons, it is not yet clear if begomovirus DNA-B sequences are more or less prone to recombination than DNA-A more broadly. The expectation is that the genomic structure of DNA-B segments (where there are no overlapping genes and a larger proportion of noncoding regions relative to the highly overlapping and mostly coding DNA-A segments) imposes fewer selective constraints on recombinants than in DNA-A sequences (44, 65) and can tolerate greater nucleotide diversity, which we do observe (Tables 3, 5). However, there are several factors that might explain why fewer recombinants were detected among the DNA-B datasets. No reliable, complete alignments were obtained due to the high divergence of DNA-B segments, hindering our ability to characterize recombination events using RDP4. RDP4 analyses were also explicitly set up to be conservative and to test for intrasegment recombination in this study, so additional events involving DNA-A and DNA-B segments might have been missed. Additionally, CMB DNA-B sequences are infrequently sampled compared to DNA-A sequences (Table 1), which reduces our ability to detect recombination both by having a lower number of exemplar parental sequences in our datasets and by having fewer representative DNA-B sequences. A recent study suggests that recombination occurs more frequently in DNA-B segments of New World begomoviruses relative to their cognate DNA-As, but this pattern may be virus-specific (66). Moreover, previous research suggests that DNA-A and DNA-B sequences have been subjected to different evolutionary pressures which have resulted in distinct evolutionary histories for the two segments, with further segment-specific differences found between New World and Old World begomoviruses (44). Regardless of the absolute rate of recombination differences that may exist between DNA-A and DNA-B segments, ρ/θ values for groups of DNA-B sequences follow the trend of DNA-A segments, which suggests that mutations occur more frequently than recombination events.

Like mutations, viral recombination events are usually deleterious (67–69). However, previous studies of begomovirus recombination have shown that there is a subset of fit recombinants that can be generated in the lab (70, 71) and be observed in nature (72). Recombination can additionally recover functional full-length genomes from populations with defective geminiviruses (69, 73, 74). Since begomovirus phenotypes associated with recombination include altered disease severity (75, 76), host range expansion (76, 77) and resistance-breaking (78, 79), recombination is also a major contributor to the epidemic potential of these viruses. Consequently, recombination is a markedly important evolutionary mechanism with epidemiological implications for begomovirus emergence. Knowledge about mechanistic patterns and selective determinants of fit CMB recombinants should be incorporated in the development of anti-viral strategies to reduce the likelihood of the emergence of virulent recombinants. This is especially important in the context of breeding CMD-resistant cassava varieties which has been the most effective approach for disease control to date (80).

### Species constructed on sequence divergence are ripe for speciation by recombination

It should be noted that recombination as a driver of speciation is also a function of the way the community defines species in *Begomovirus.* Current taxonomy guidelines state that a begomovirus species is defined as a group of DNA-A isolates sharing ≥91% pairwise nucleotide sequence identity and any new isolate is assigned to a species if it shares at least 91% nt sequence identity to any one isolate from that species (81). As we increase surveying efforts of natural CMB biodiversity with improving sequencing techniques, a larger fraction of the tolerated sequence diversity within each species will be found. It will be increasingly unlikely that sufficient mutations to create >9% sequence diversity will accumulate quickly enough without any intermediates being sampled. These cataloged intermediates then shift the goal posts for “speciation,” as a novel species would have to have >9% sequence divergence from them. Consequently, recombination may be the main way to obtain enough genetic variation to cross the species demarcation threshold for begomoviruses and is therefore the likely predominant mechanism of speciation for the entire genus.

Virus speciation is often discussed in terms of ecological factors, where host specificities and virus-host interactions lead to the evolution of diverged lineages that develop into different viral species (82–85). Under this model, frequent recombination homogenizes viral diversity, and only when recombination is limited do lineages diversify (86, 87). Here, by zooming in to the CMD species complex, we provide evidence that diversity at the species level can be predominantly shaped by recombination as well. Recombination has also been implicated as a direct mechanism of speciation in other virus groups, e.g., *Luteovirideae* (88), *Bromoviridae* (89), *Reoviridae* (90) and *Papillomaviridae* (91). Additionally, recombination has shaped some deep phylogenetic relationships among viruses. Within *Geminiviridae*, most genera have emerged from ancient inter-generic recombination events (92–97). For higher taxonomic ranks, it is apparent that the origins of multiple families within *Cressdnaviricota*, including *Geminiviridae* (98, 99), can be traced to independent recombination events involving prokaryotic plasmids and diverse plant and animal RNA viruses (100). These and other recombination events in deep phylogeny have led to both modular patterns of virus evolution and polyphyletic groupings across the Baltimore classifications (101).

The general trend of speciation via recombination for CMBs might not be true for species in other virus families. For instance, a recent review of potyviruses found recombination to be common within populations but uncommon as a mechanism of speciation (102). In picornaviruses, whose species demarcation is defined by a significant degree of amino acid identity, it has been concluded that recombination limits speciation and members of distinct species based on current taxonomic schemes are so diverged that they are generally presumed to be incompatible (103). Our contrasting results are likely due to the narrow way that novel species are determined in begomoviruses (<91% nucleotide identity for the DNA-A segment). However, as more viral groups move to nucleotide identity as species demarcation criteria as a way to integrate the wealth of viral diversity known from genetic sequences alone (104), our conclusions from CMBs may prove more broadly applicable.

## MATERIALS AND METHODS

### CMB sequence data sets

Two data sets comprised of all full-length DNA-A and DNA-B nucleotide sequences corresponding to the 11 recognized CMB species were downloaded from the GenBank database via NCBI Taxonomy Browser (https://www.ncbi.nlm.nih.gov/Taxonomy/Browser/wwwtax.cgi) between January and July 2019. The 11 species analyzed here (and corresponding virus abbreviations) are *African cassava mosaic virus* (ACMV), *African cassava mosaic Burkina Faso virus* (ACMBFV), *Cassava mosaic Madagascar virus* (CMMGV), *South African cassava mosaic virus* (SACMV), *East African cassava mosaic virus* (EACMV), *East African cassava Cameroon virus* (EACMCV), *East African cassava Kenya virus* (EACMKV), *East African cassava Malawi virus* (EACMMV), *East African cassava mosaic Zanzibar virus* (EACMZV), *Sri Lankan cassava mosaic virus* (SLCMV) and *Indian cassava mosaic virus* (ICMV). All sequences were organized to begin at the nick site of the conserved nonanucleotide motif at the origin of replication (5’-TAATATT//AC-3’).

### Alignments and sample classification

All multiple sequence alignments were constructed using the MUSCLE method (105) as implemented in MEGA X (106) and manually corrected using AliView v1.26 (107). Multiple alignments have been archived as Zenodo records: 11 species DNA-A alignment (https://zenodo.org/record/4029589), CMMGV, EACMCV, EACMV, EACMKV, EACMMV, EACMZV, SACMV DNA-B alignment (https://zenodo.org/record/3965023), ACMV and ACMBFV DNA-B alignment (https://zenodo.org/record/3964979), and ICMV and SLCMV DNA-B (https://zenodo.org/record/3964977).

A pairwise nucleotide identity matrix was calculated for complete DNA-A sequences using SDT v1.2 (108) and was used to assign each DNA-A sequence to a viral species according to the ICTV-approved begomovirus species demarcation threshold of >91% DNA-A identity (81). For DNA-B sequences, the species assignment listed in GenBank was used; species definitions for DNA-B are less distinct, as discussed in the text.

### Phylogenetic network analysis

Phylogenetic networks, which can capture conflicting phylogenetic signals such as those caused by recombination, were inferred from the alignments using the distance-based Neighbor-Net method (109) implemented in SplitsTree4 v4.14 (110). Distances were corrected with a GTR + G model of sequence evolution using base frequencies, rate heterogeneity and gamma shape parameters estimated with jModelTest v2.1.6 (111) on XSEDE.

### Recombination detection and similarity plots

Putative recombinants, and major and minor “parents” within the data sets were determined using the RDP (112), GeneConv (11), Bootscan (113), MaxChi (114), Chimaera (115), SiScan (116), and 3Seq (117) recombination detection methods implemented on the RDP4 v4.100 suite (118). The terms ‘major parent’ and ‘minor parent’ are used by RDP4 to refer to sequences that have respectively contributed the larger and smaller fractions to the recombinant and are regarded as closest relatives to the true isolates involved in the event. Analyses were performed with default settings, while also enabling Chimaera and 3Seq for primary scan, and a Bonferroni-corrected P-value cutoff of 0.05 was used. Only events supported by at least five of the seven methods were considered high-confidence events. Breakpoint positions, putative recombinants and “parental” sequences were evaluated and manually adjusted when necessary using the available breakpoint cross-checking tools and phylogenetic tree construction methods available in RDP4. RDP4 results files have been archived as Zenodo records: RDP4 results for ACMBFV, ACMV, CMMGV, EACMCV, EACMV, EACMKV, EACMMV, EACMZV, SACMV DNA-A (https://zenodo.org/record/4592854), RDP4 results for ICMV and SLCMV DNA-A (https://zenodo.org/record/4592926), RDP4 results for EACMV-like + CMMGV DNA-B (7 “species”) (https://zenodo.org/record/3965029), RDP4 results for ACMV and ACMBFV DNA-B (https://zenodo.org/record/3975834), and (RDP4 results for ICMV and SLCMV DNA-B (https://zenodo.org/record/3975838). Events were considered as macroevolutionary recombination events when all members of a designated species had evidence of said event. A BLASTn analysis of the ‘non-redundant nucleotide’ database in NCBI (https://blast.ncbi.nlm.nih.gov/Blast.cgi) was performed to identify the species whose members have sequences most similar to the “Unknown” recombinant fragments in our data sets.

Based on RDP4 results, similarity analyses comparing the recombinants to their putative parental sequences were performed using SimPlot v3.5.1 (119). All plots were done using the Kimura 2-parameter distance model with a sliding window size of 100 and a step size of 10. For the the similarity plots in Figure 2, analyses were done by comparing 50% consensus sequences of all members of the compared species, respectively (except in the case of 2C where the best candidate sequence was used). Similarity plots for events in Tables 2 and 4 were made comparing the best candidate recombinant and parent sequences reported by RDP4 and are presented in the appendix.

### Estimates for the relative rates of recombination and mutation, linkage disequilibrium correlations with distance and likelihood permutation tests of recombination

LDhat v2.2 (43) was used to infer composite likelihood estimates (CLEs) of population-scaled recombination rates (*ρ* = 2*N*_*e*_*r*) and estimates of population-scaled mutation rates (*θ*^*w*^ = 2*N*_*e*_*μ*) with the PAIRWISE and CONVERT packages, respectively. This program uses an extension of Hudson’s composite-likelihood method (120), which estimates the population recombination rate by combining the coalescent likelihoods of all pairwise comparisons of segregating sites. The extension in LDhat allows for a finite-sites mutation model, which makes it appropriate for sets of sequences with high mutation rates such as the ones found in viral genomes.

CONVERT was used with all default settings. While using PAIRWISE, a gene-conversion model with an average tract length of 500 nt was fitted, and a precomputed likelihood lookup table for per-site θ=0.01 with a maximum 2*N*_*e*_*r* of 100 and 101-point size grid was used to obtain the CLEs of *ρ*. Since precomputed likelihood lookup tables for data sets larger than n=100 are not available, the COMPLETE package was used with GNU parallel (https://zenodo.org/record/3903853) to generate a likelihood lookup table for per-site θ=0.01 that can accommodate data sets of up to 320 sequences to use for the larger data sets in this study. File available as “LDhat coalescent likelihood lookup table for 320 sequences with theta = 0.01” (https://zenodo.org/record/3934350). *ρ*/*θ*^*w*^ estimate was obtained as the relative rate of recombination and mutation in the history of the samples within each analyzed clade. Additionally, PAIRWISE was used to obtain the correlation between estimates of linkage disequilibrium (r^2^) and physical distance (d), and to test for the presence of recombination using the likelihood permutation test (LPT) developed by McVean et al. (43).

### Standing genetic diversity and divergence between parental and recombinant species

The per-site standing genetic diversity of each species was assessed by calculating nucleotide diversity π (121), which is the average number of pairwise nucleotide differences per site between sequences within a clade. To obtain absolute measures of divergence between recombinant and parental species, per-site D_XY_ (121) was calculated. D_XY_ refers to the average number of pairwise differences between sequences from two clades while excluding all intra-clade comparisons and is calculated as:

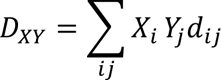

where, in two clades, X and Y, *dij* measures the number of nucleotide differences between the i^th^ haplotype from X and the j^th^ haplotype from Y. All per-site estimates were obtained with DnaSP v6.12 (122). When estimating nucleotide diversity, gaps/missing information were excluded only in pairwise comparisons. For sliding window analyses, a sliding window size of 100 nt (including gaps) and a step size of 10 nt were used.

### Data availability

As noted above, multiple alignments, RDP4 results and the generated LDhat likelihood lookup table have been deposited as Zenodo records. Alignment for DNA-A: 11 species DNA-A alignment (https://zenodo.org/record/4029589), Alignments for DNA-B: CMMGV, EACMCV, EACMV, EACMKV, EACMMV, EACMZV, SACMV DNA-B alignment (https://zenodo.org/record/3965023) ACMV and ACMBFV DNA-B alignment (https://zenodo.org/record/3964979) ICMV and SLCMV DNA-B (https://zenodo.org/record/3964977) RDP4 results for DNA-A: RDP4 results for ACMBFV, ACMV, CMMGV, EACMCV, EACMV, EACMKV, EACMMV, EACMZV, SACMV DNA-A (https://zenodo.org/record/4592854) RDP4 results for ICMV and SLCMV DNA-A (https://zenodo.org/record/4592926) RDP4 results for DNA-B: RDP4 results for EACMV-like + CMMGV DNA-B (7 “species”) (https://zenodo.org/record/3965029) RDP4 results for ACMV and ACMBFV DNA-B (https://zenodo.org/record/3975834) (RDP4 results for ICMV and SLCMV DNA-B (https://zenodo.org/record/3975838) LDhat coalescent likelihood lookup table for 320 sequences with theta = 0.01 (https://zenodo.org/record/3934350)

## ACKNOWLEDGEMENTS

We are grateful to all the research groups that have shared sequence data via GenBank and the farmers who enabled collection of that data. We would like to thank members of the Duffy lab at Rutgers University and our colleagues at the Hanley-Bowdoin lab in North Carolina State University for feedback and a critical reading of this manuscript. We also thank the staff of the Office of Advanced Research Computing (OARC) at Rutgers for access to and maintenance of the Amarel cluster. This work was supported by US NSF award OIA-1545553 to SD and an HHMI Gilliam Fellowship for Advanced Study for ACB. DD was partly supported by the Aresty Undergraduate Research Scholars Program at Rutgers University.

## APPENDIX FIGURE LEGENDS

**Appendix:** Distance plots and schematic diagrams for all high-confidence recombination events. The distance plots that follow confirm the plausibility of the RDP4-identified recombination events listed in Tables 2 and 4. Note that Kimura-2-parameter-model-corrected similarities calculated by SimPlot are distinct from percent nucleotide identity but are presented on the conventional scale.

Figure A1. **Similarity plots for four African CMB DNA-A macroevolutionary recombination events.** Note that BLAST analysis suggests that a ToLCCMV-like was a more likely donor for Event 1 (Figure 1) but similarity for the closest sequence in the dataset (CMMGV) is plotted.

Figure A2. **Similarity plot for EACMKV DNA-A macroevolutionary recombination event, exemplifed by AJ717578.** Note that the y-axis limits differ from other graphs.

Figure A3. **Similarity plots for 3 African CMB DNA-A recombinant haplotypes.** Note that the y-axis limits differ from other graphs. Comparisons correspond to RDP4 results for events 7-9.

Figure A4. **Similarity plots for 4 African CMB DNA-A recombination events.** Comparisons correspond to RDP4 results for events 10-13. The color used to represent EACMV sequences in event 11 is slightly different from other graphs to increase color contrast.

Figure A5. **Similarity plots for 4 African CMB DNA-A recombination events.** Comparisons correspond to RDP4 results for events 14-17. Note that event 16 corresponds to the KE2 recombinant haplotype described in Bull et al. (31). The color used to represent EACMV sequences in event 16 is slightly different from other graphs to increase color contrast.

Figure A6. **Similarity plots for 4 Asian CMB DNA-A recombination events.** Comparisons correspond to RDP4 results for macroevolutionary event 6 and events 18-20.

Figure A7. Similarity plots for 4 Asian CMB DNA-A recombination events. Comparisons correspond to RDP4 results for events 21-24.

Figure A8. **Similarity plot for ACMBFV DNA-B putative recombination event and sequence alignments depicting potential recombination scenarios.** Note that the sole available ACMBFV DNA-B sequence did not meet our criterion for a high-confidence recombination event, as described above, but is highlighted for comparison to SLCMV DNA-B. The putative Rep protein binding site of the ACMBFV DNA-B isolate includes a single core Rep (AC1) binding sequence (GGGGT, highlighted in blue) with a potential inverted repeat (GGACC, highlighted in green). Scenario 1 shows the possibility that ACMBFV DNA-B originated from a recombination event involving an ACMBFV DNA-A that donated part of the CR to an ACMV DNA-B, which has a distinct binding site (27) (GGAGA, highlighted in pink). Scenario 2 shows the possibility that ACMBFV DNA-B originated from a recombination event involving a different virus segment and ACMV DNA-B. Based on best BLAST hit, the minor parent for Scenario 2 could be a relative of a tomato leaf curl Nigeria virus (ToLCNGV) segment (accession: FJ685621). The single Rep-binding sequence and inverted repeat for ToLCNGV match the ones for ACMBFV DNA-B and have been characterized previously (123). The C1 ORF TATA box for each sequence is shown in a box. Grey sites in the aligned sequences correspond to sites that are distinct from the ACMBFV DNA-B sequence.

Figure A9. **Similarity plots for 4 EACMV-like DNA-B recombination events.** Comparisons correspond to RDP4 results for clade forming event B1 and events B3, B7-B8.

Figure A10. **Similarity plots for 3 EACMV-like DNA-B recombination events resulting from relatively similar sequences.** Comparisons correspond to RDP4 results for events B4-B6.

Figure A11. **Similarity plots for 3 Asian CMB DNA-B events.** Comparisons correspond to RDP4 results for clade-forming event B2 and events B9-B10. Crespo-Bellido et al.

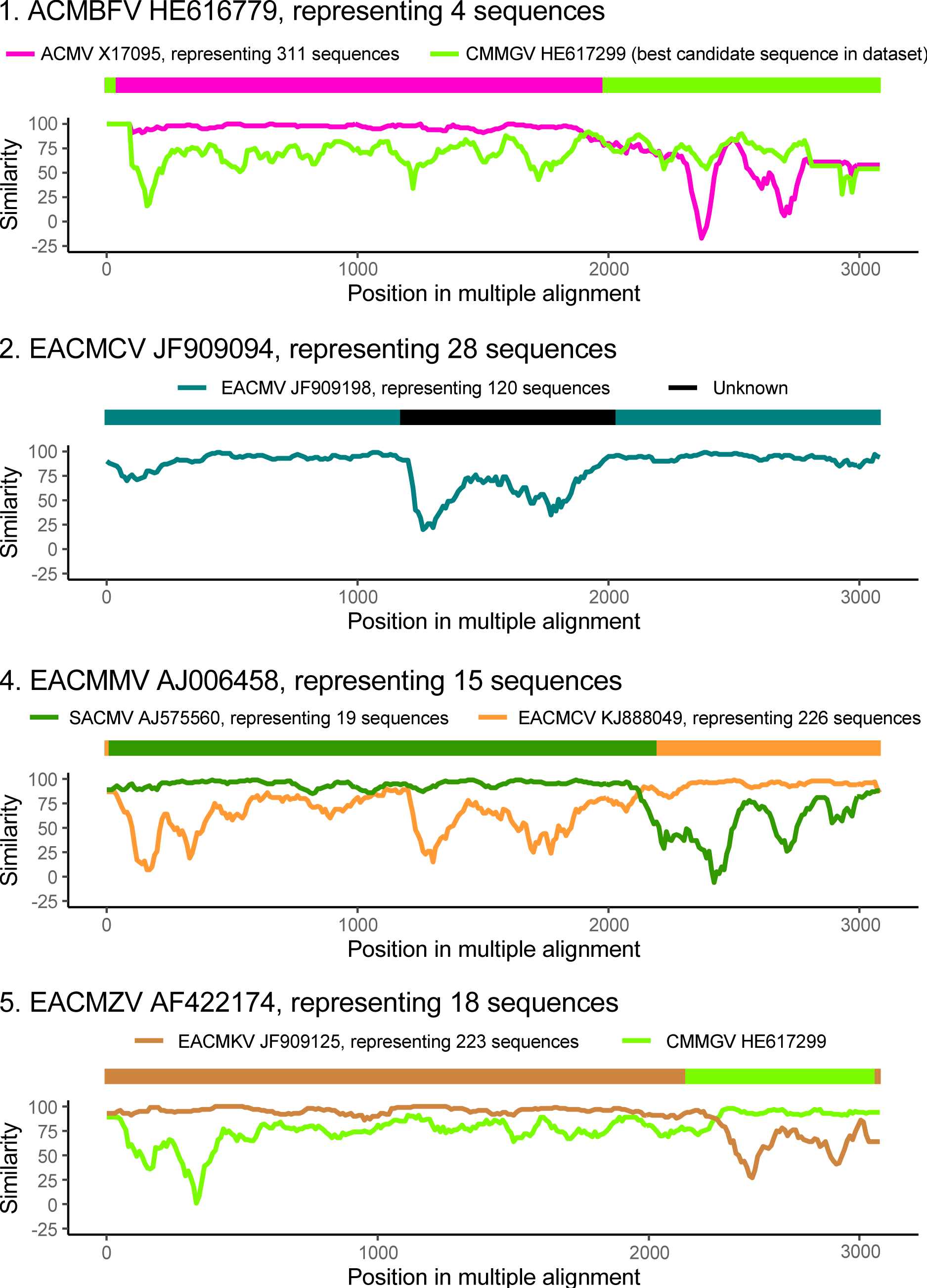
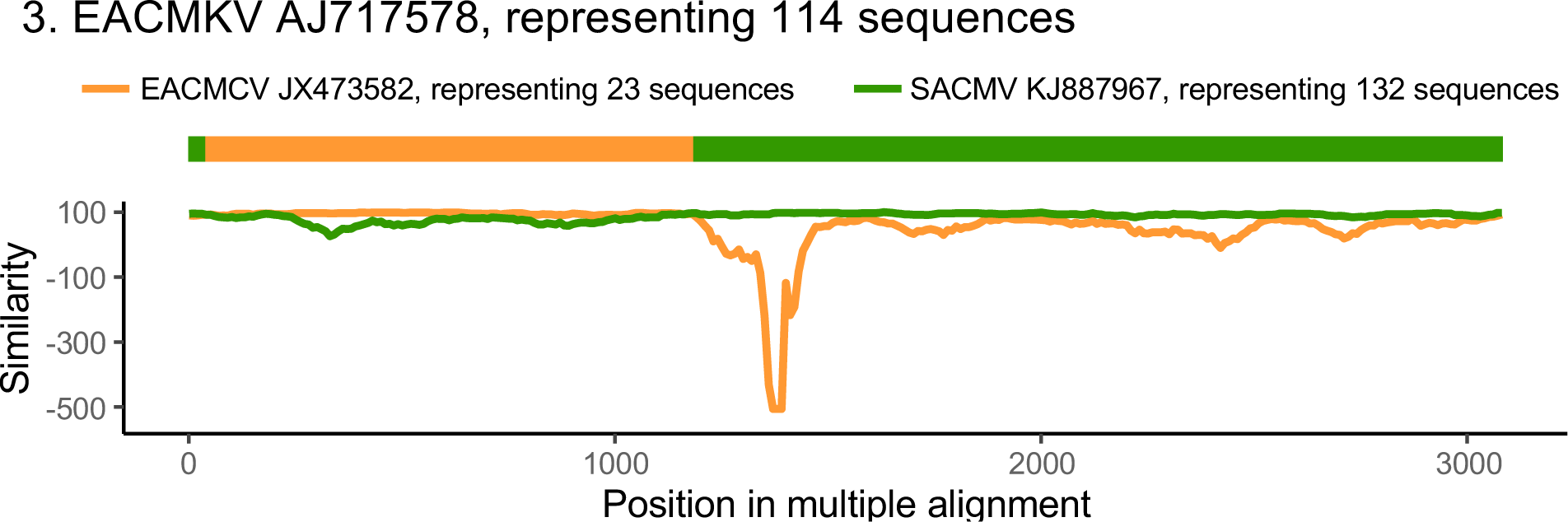
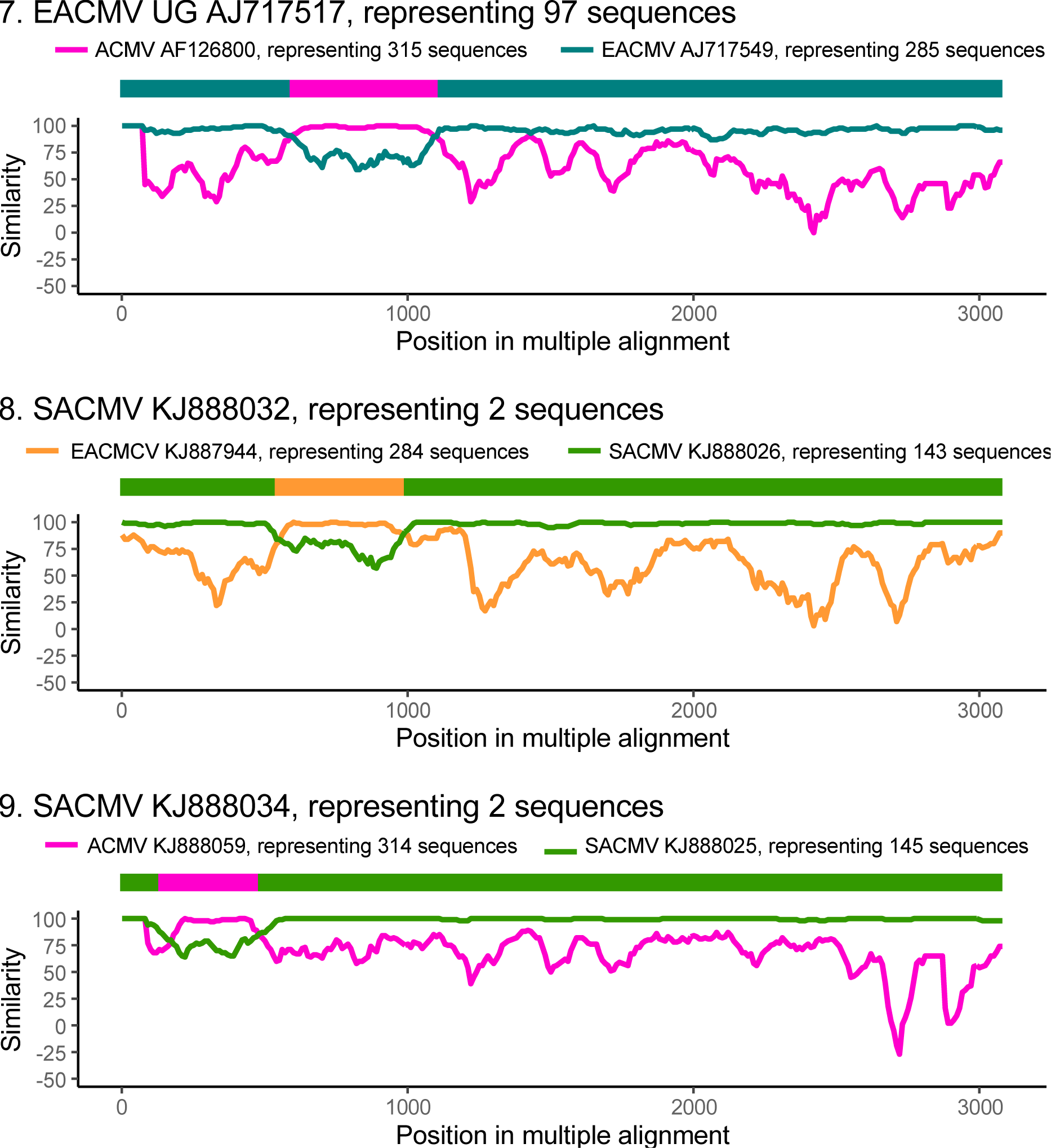
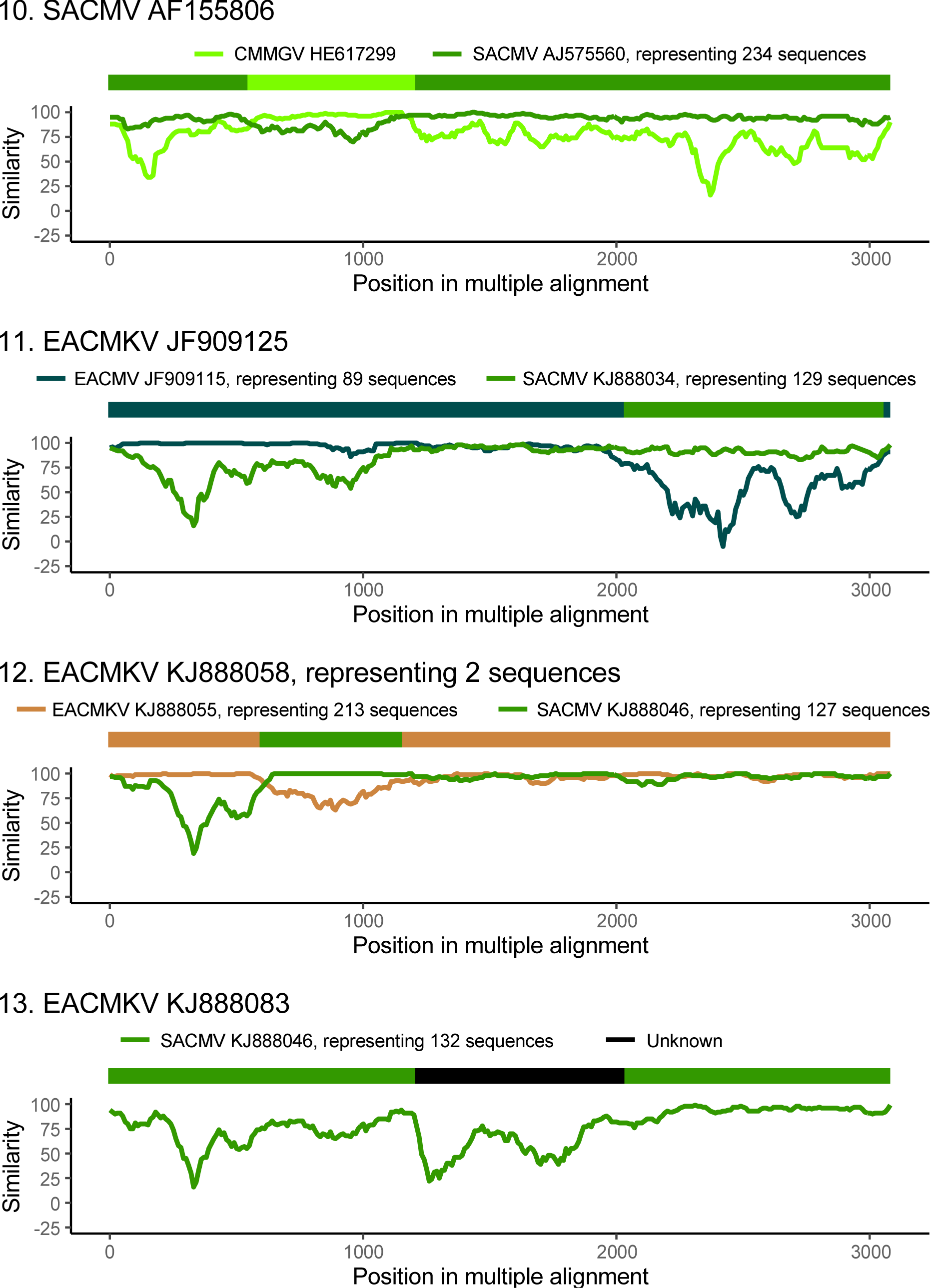
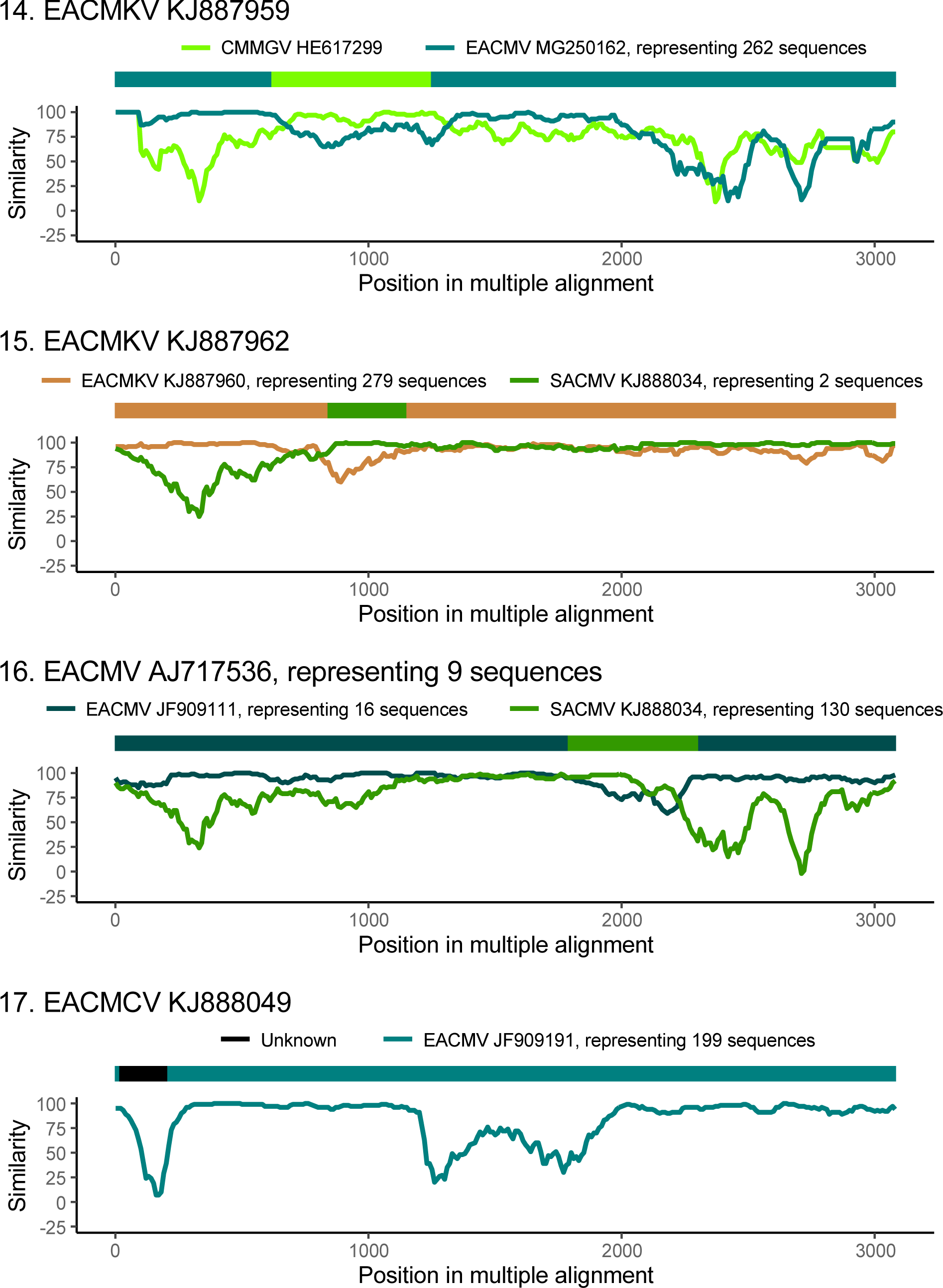
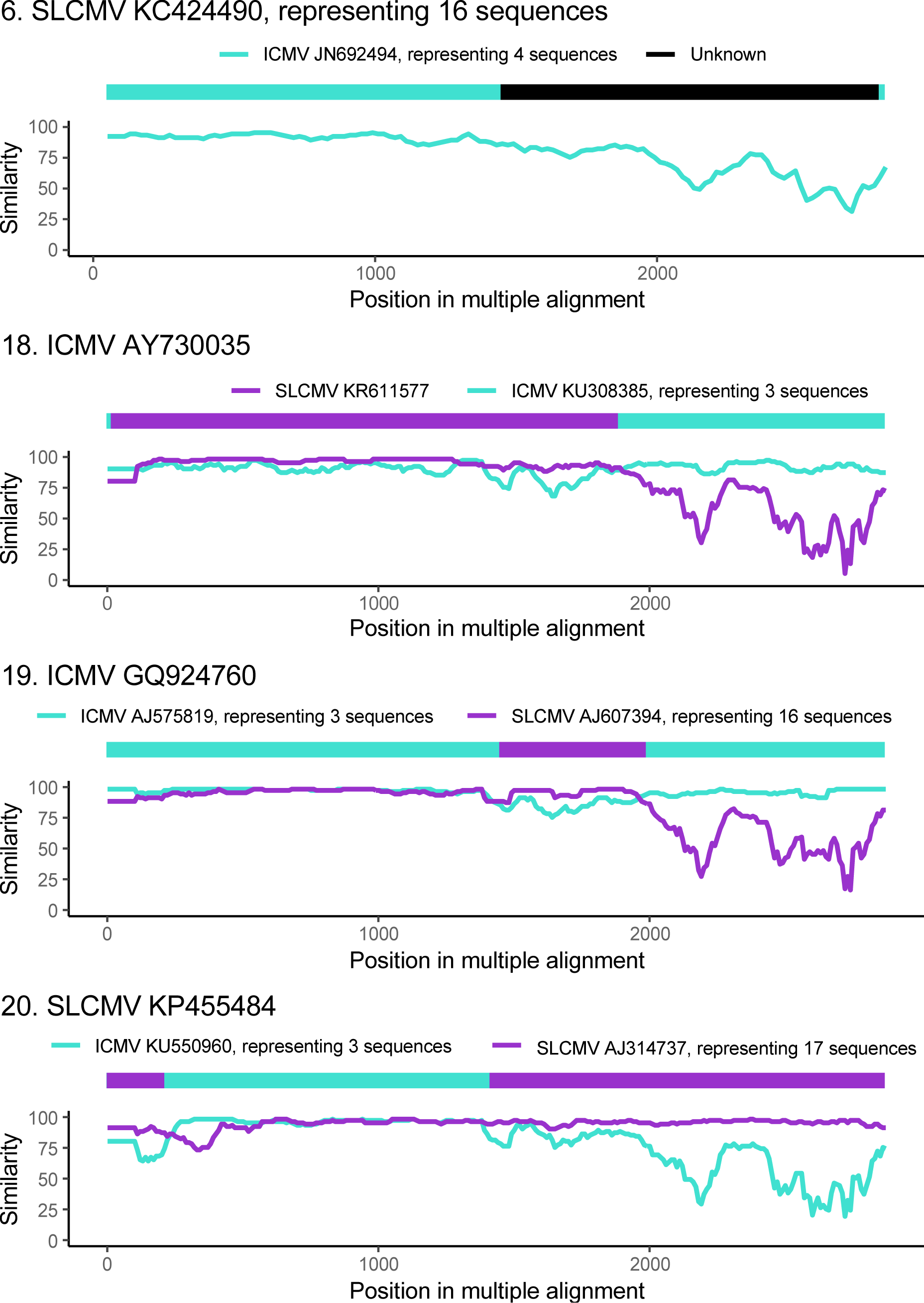
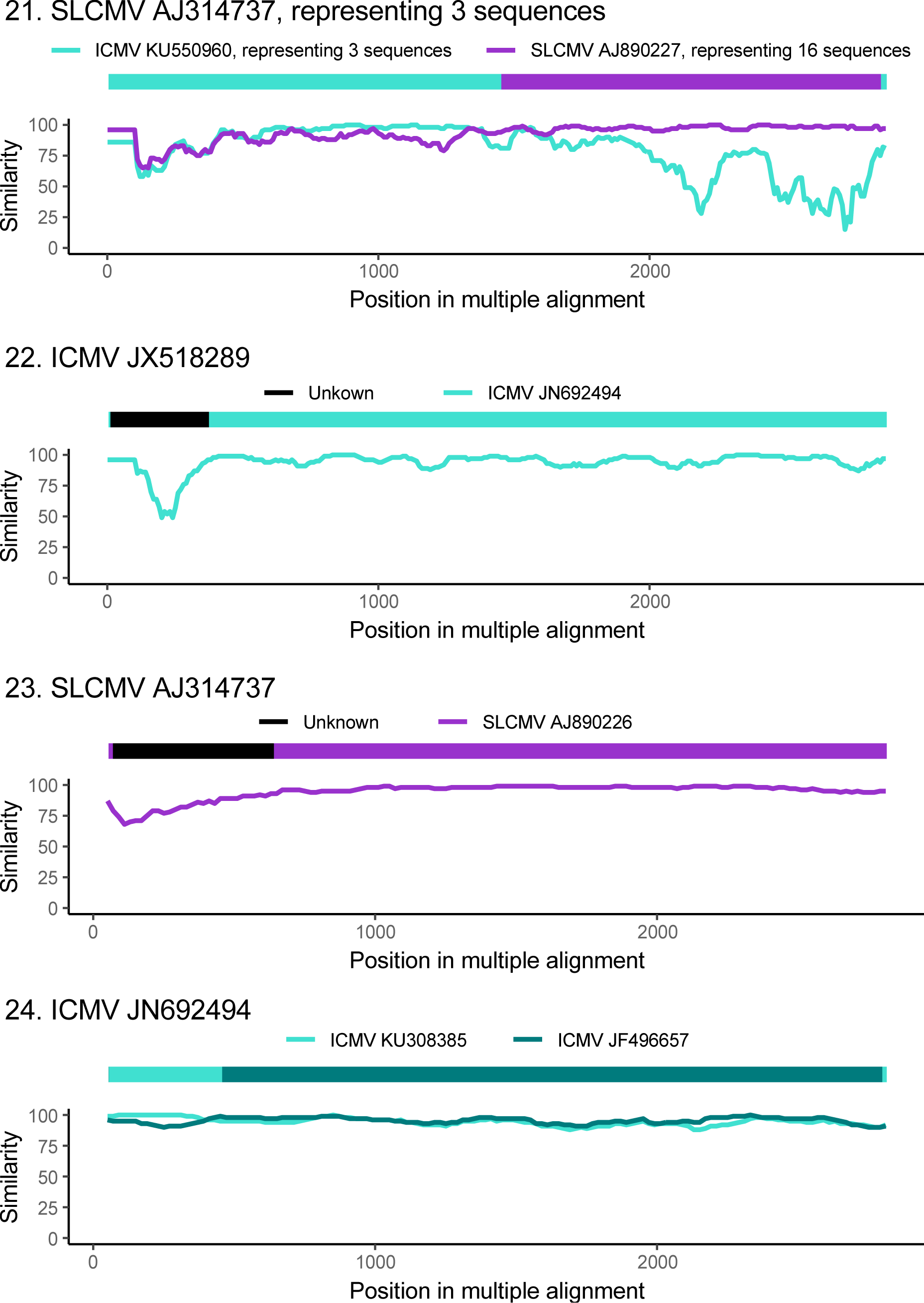
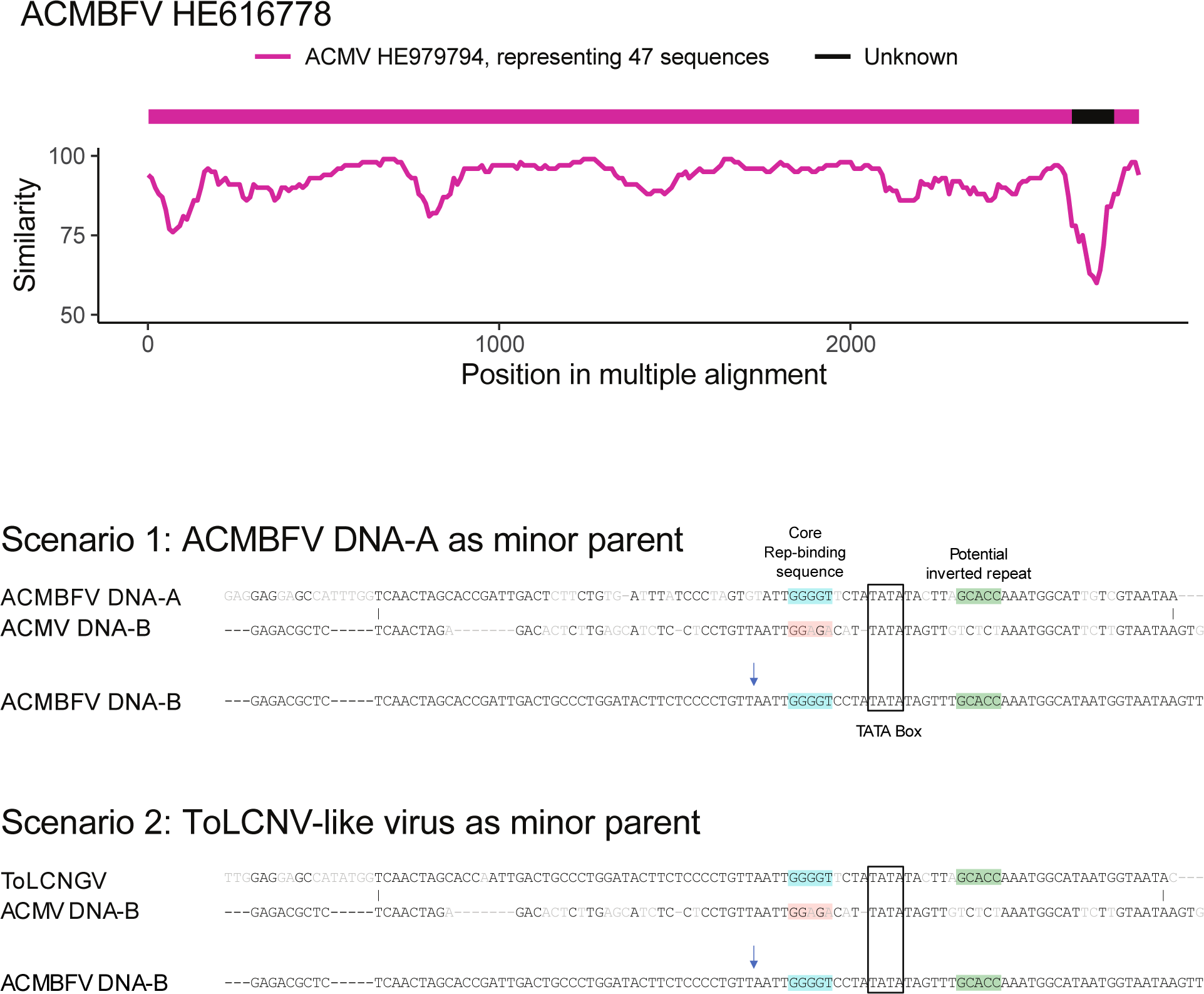
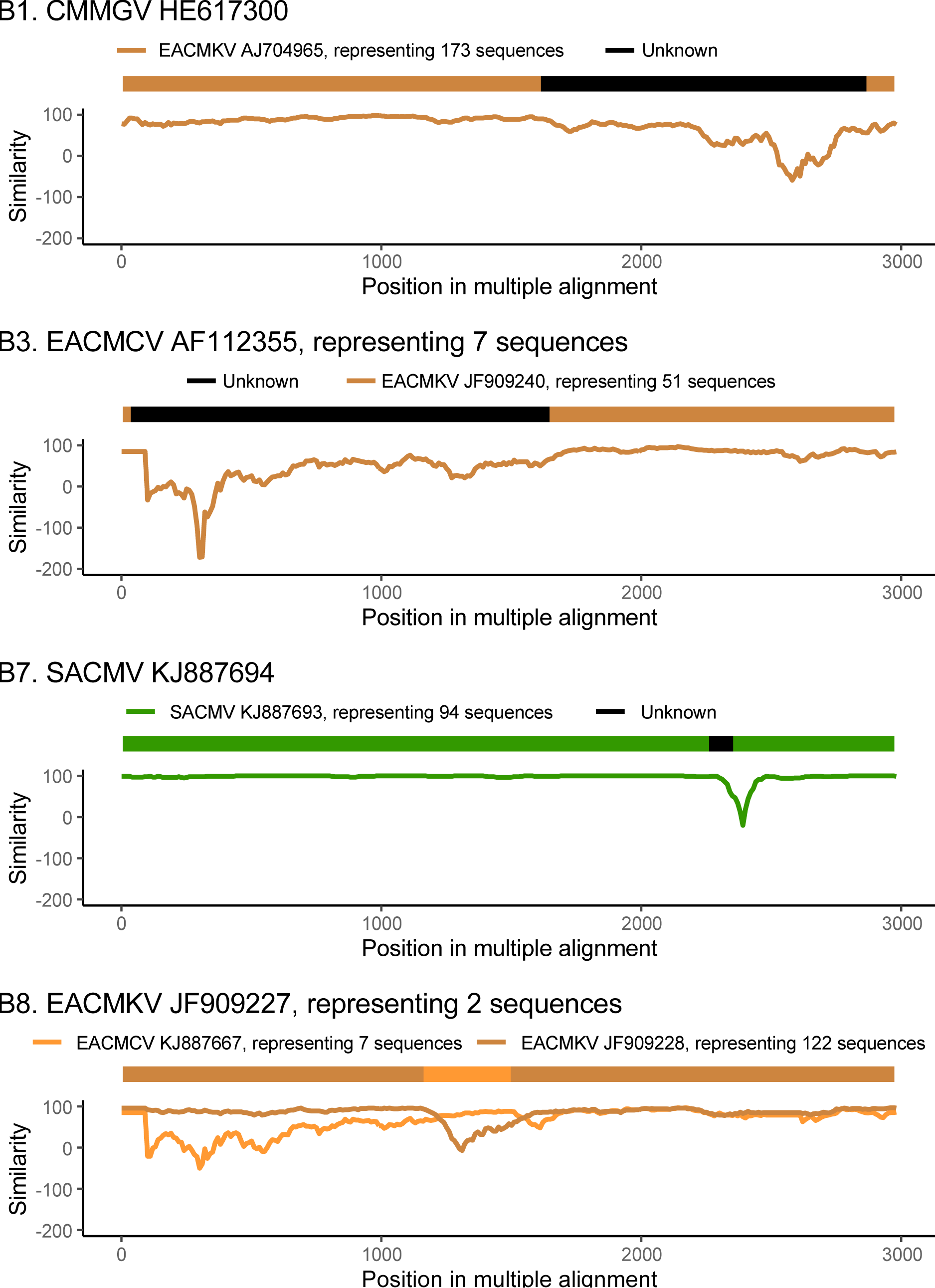
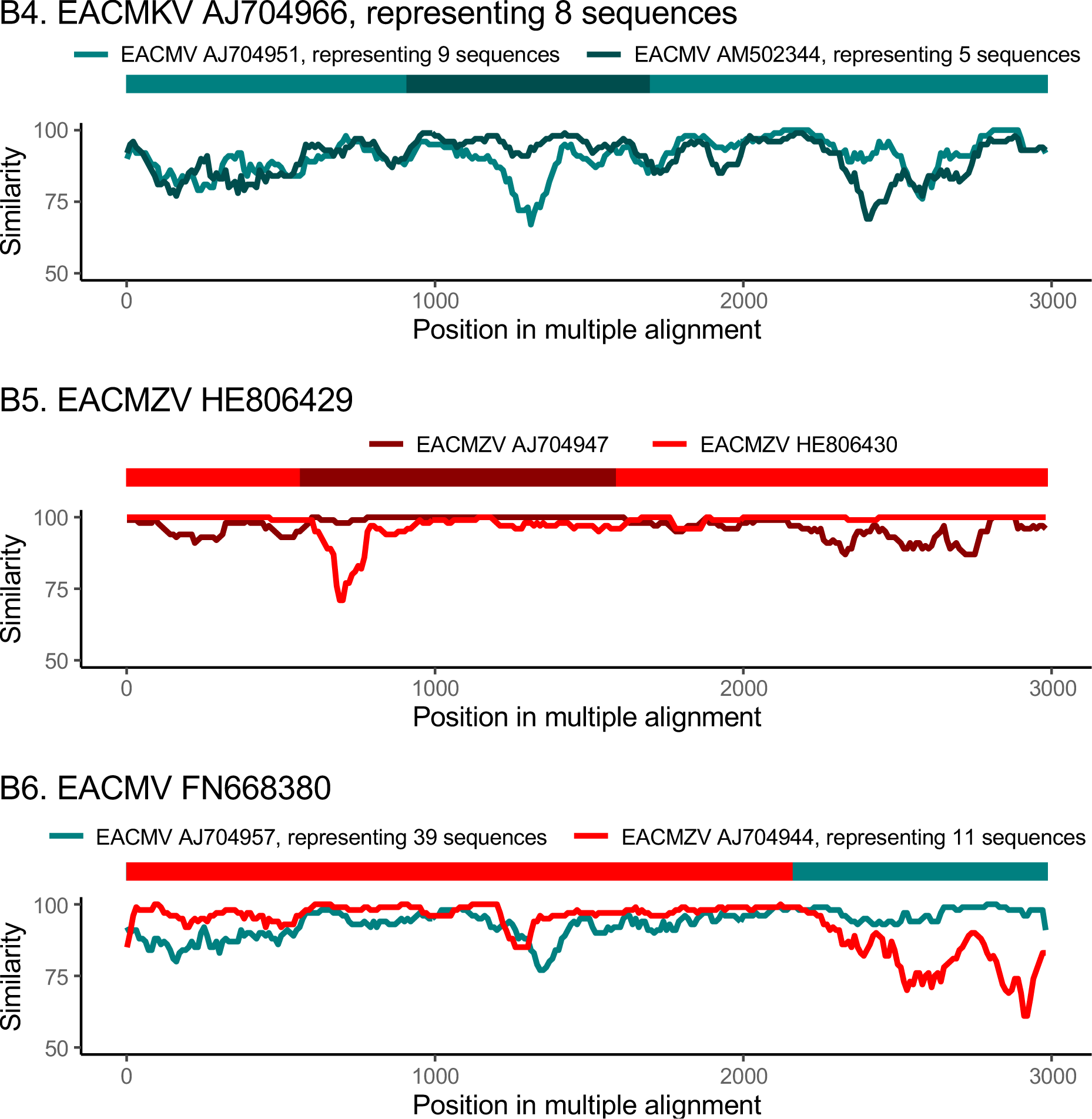
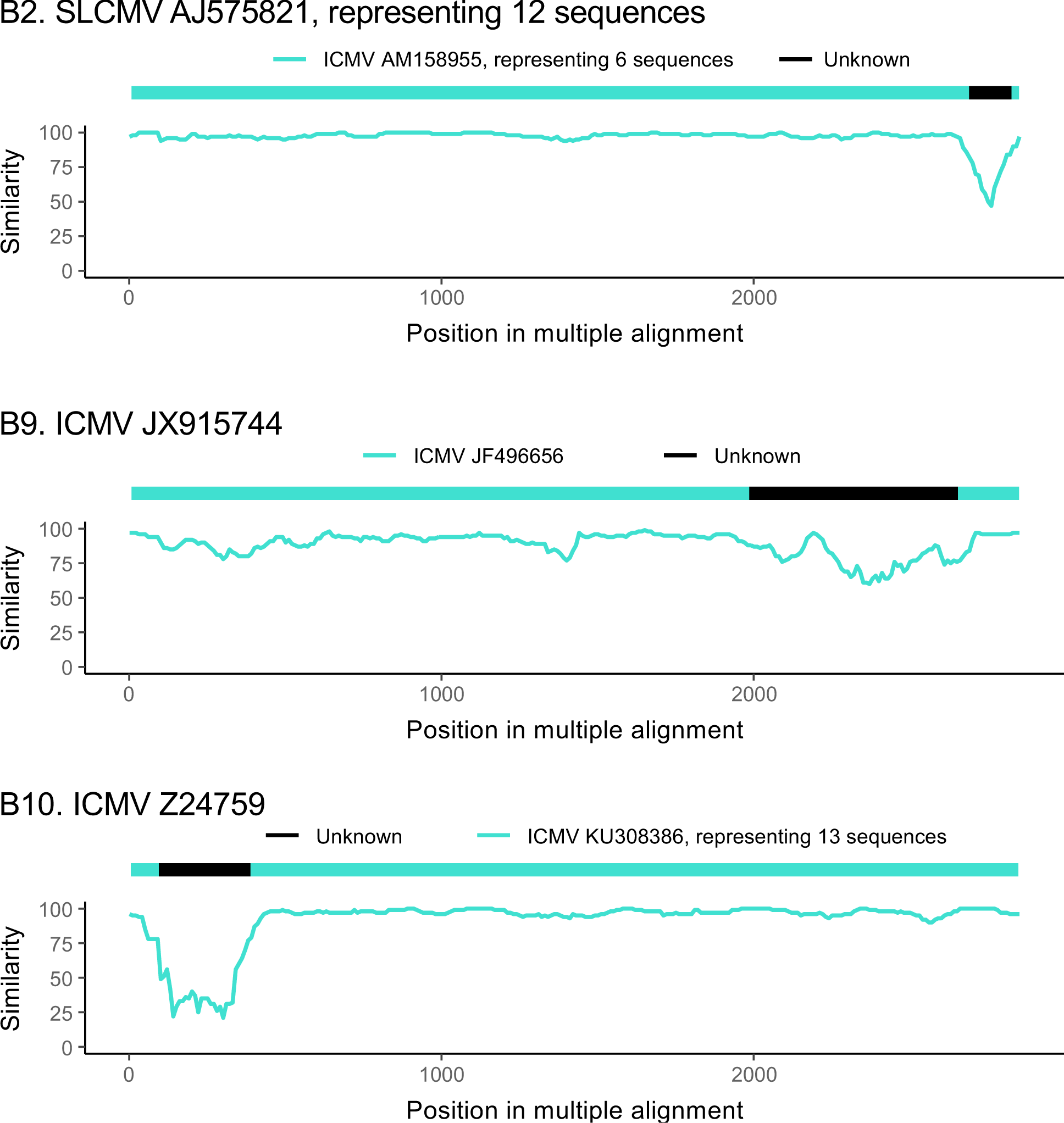

